# Mapping transcriptional responses to cellular perturbation dictionaries with RNA fingerprinting

**DOI:** 10.1101/2025.09.19.676866

**Authors:** Isabella N. Grabski, Junsuk Lee, John Blair, Carol Dalgarno, Isabella Mascio, Alexandra Bradu, David A. Knowles, Rahul Satija

## Abstract

Single-cell perturbation dictionaries provide systematic measurements of how cells respond to genetic and chemical perturbations, and create the opportunity to assign causal interpretations to observational data. Here, we introduce RNA fingerprinting, a statistical framework that maps transcriptional responses from new experiments onto reference perturbation dictionaries. RNA fingerprinting learns denoised perturbation “fingerprints” from single-cell data, then probabilistically assigns query cells to one or more candidate perturbations while accounting for uncertainty. We benchmark our method across ground-truth datasets, demonstrating accurate assignments at single-cell resolution, scalability to genome-wide screens, and the ability to resolve combinatorial perturbations. We demonstrate its broad utility across diverse biological settings: identifying context-specific regulators of p53 under ribosomal stress, characterizing drug mechanisms of action and dose-dependent off-target effects, and uncovering cytokine-driven B cell heterogeneity during secondary influenza infection in vivo. Together, these results establish RNA fingerprinting as a versatile framework for interpreting single-cell datasets by linking cellular states to the underlying perturbations which generated them.

## INTRODUCTION

Despite dramatic advances in single-cell and spatial profiling technologies^1–10^, a fundamental challenge remains: most genomic measurements are observational in nature. Even the most comprehensive transcriptional atlases, which reveal rich variation in cell types, states, and trajectories between healthy and diseased samples^11–15^, often fall short in determining underlying mechanisms. While such transcriptional signatures can clearly describe how cellular states vary across conditions, they offer limited information about why those differences arise, and disentangling these upstream causes is inherently difficult. As a result, despite their immense descriptive power, most genomic datasets remain correlative, making causal inference a persistent and central challenge in the field^16^.

In contrast, perturbation experiments provide direct causal insight^9,17,18^. When a gene is perturbed or a cell is exposed to a defined stimulus, the resulting transcriptional response represents a “fingerprint” that encodes the causal consequences of the perturbation. The recent proliferation of single-cell perturbation dictionaries – datasets that systematically record cellular responses to an array of genetic, chemical, or immunological perturbations – provide a powerful new opportunity to assign causal mechanisms in single-cell data. These dictionaries include genome-scale CRISPRi screens^19^, drug response atlases like sci-Plex^20^ and Tahoe 100M^21^, and immune stimulation datasets profiling thousands of cytokine-cell type combinations in vivo^22^.

As these resources grow in scale and diversity, they can serve as references for interpreting new data. For example, a match between the transcriptional profile of a drug-treated cell and that of a known gene perturbation provides mechanistic insight into the drug’s target or pathway of action^23^. More broadly, any new experimental condition, such as a disease state or environmental exposure, could in principle be mapped to a perturbation dictionary to infer potential causal factors. This concept of matching across datasets to learn underlying molecular mechanisms has been explored extensively using bulk transcriptomic measurements, particularly in the drug characterization context^23–25^. For instance, one recent analysis found that the top 100 most correlated perturbations to a query signature were enriched for the expected target^26^, but the ability to make more precise matches has remained challenging. In contrast to bulk measurements, single-cell datasets enable highly multiplexed profiling with minimal batch effects and allow for the modeling of heterogeneous cellular responses, which we propose can boost the power and specificity of perturbation matching.

Despite the potential of emerging single-cell dictionaries, currently available single-cell computational frameworks are not designed for this task. Existing single-cell integration and annotation algorithms rely on broad transcriptional differences across a cell’s global expression profile and assume a one-to-one correspondence between a single cell and a single label^27^. In contrast, perturbation responses are often subtle and may represent only a small component of a cell’s transcriptomic profile, can arise from combinations of multiple perturbations, and may be heterogeneous even within cells of the same subtype^28,29^. Moreover, perturbations with related functions, such as members of the same complex, often yield nearly indistinguishable phenotypes^19^, requiring new methods that can handle substantial uncertainty in matching and assignment.

Here, we introduce RNA fingerprinting, a novel statistical framework for mapping transcriptional responses from new experiments to known perturbations in reference dictionaries. Our method is compatible with a wide variety of perturbation dictionaries (“references”), and learns denoised latent representations (“fingerprints”) of each perturbation using a multi-condition factor model. To interpret new data (“queries”), we map these query data to one or more of these fingerprints using a regression approach adapted from statistical genetics^30^, summarizing complex patterns of uncertainty with Bayesian credible sets. Using ground-truth datasets, we demonstrate that RNA fingerprinting robustly and accurately infers the causal regulators driving both individual and combinatorial perturbation responses in query cells.

We apply RNA fingerprinting across a range of biological settings to uncover new mechanistic insights that would be difficult to detect using conventional approaches. By mapping perturbation responses across distinct genetic contexts, we identify RPL10 and RPL24 as previously unrecognized potential regulators of p53 activation under ribosomal impairment. In drug-treated cells, our method reveals dose-dependent off-target effects in putatively selective cancer therapeutics, including HDAC6 inhibitors whose transcriptional profiles closely resemble a fingerprint of pan-HDAC inhibition. Finally, by analyzing in vivo immune responses in influenza-rechallenged mice, RNA fingerprinting identifies a heterogeneous B cell cytokine response and nominates causal cytokines, highlighting subtle and variable immune behaviors that escape detection in standard analyses. Together, these findings demonstrate how perturbation dictionaries can serve as causal references for interpreting new single-cell data.

## RESULTS

### Mapping cellular responses with RNA fingerprinting

RNA fingerprinting is designed to infer potential causal regulators from new single-cell datasets by leveraging large-scale perturbation dictionaries as references (Figure 1A). A dictionary consists of single-cell RNA-sequencing profiles for cells that were exposed to at least one of a set of possible perturbations, as well as control (i.e. non-targeting) cells. Our framework is compatible with diverse perturbations, including genetic knockdowns, activations, drug treatments, or cytokine stimulations. In each case, we propose that these references encode a collection of transcriptional “fingerprints” that reflect the consequence of each perturbation on gene expression.

**Figure 1.**
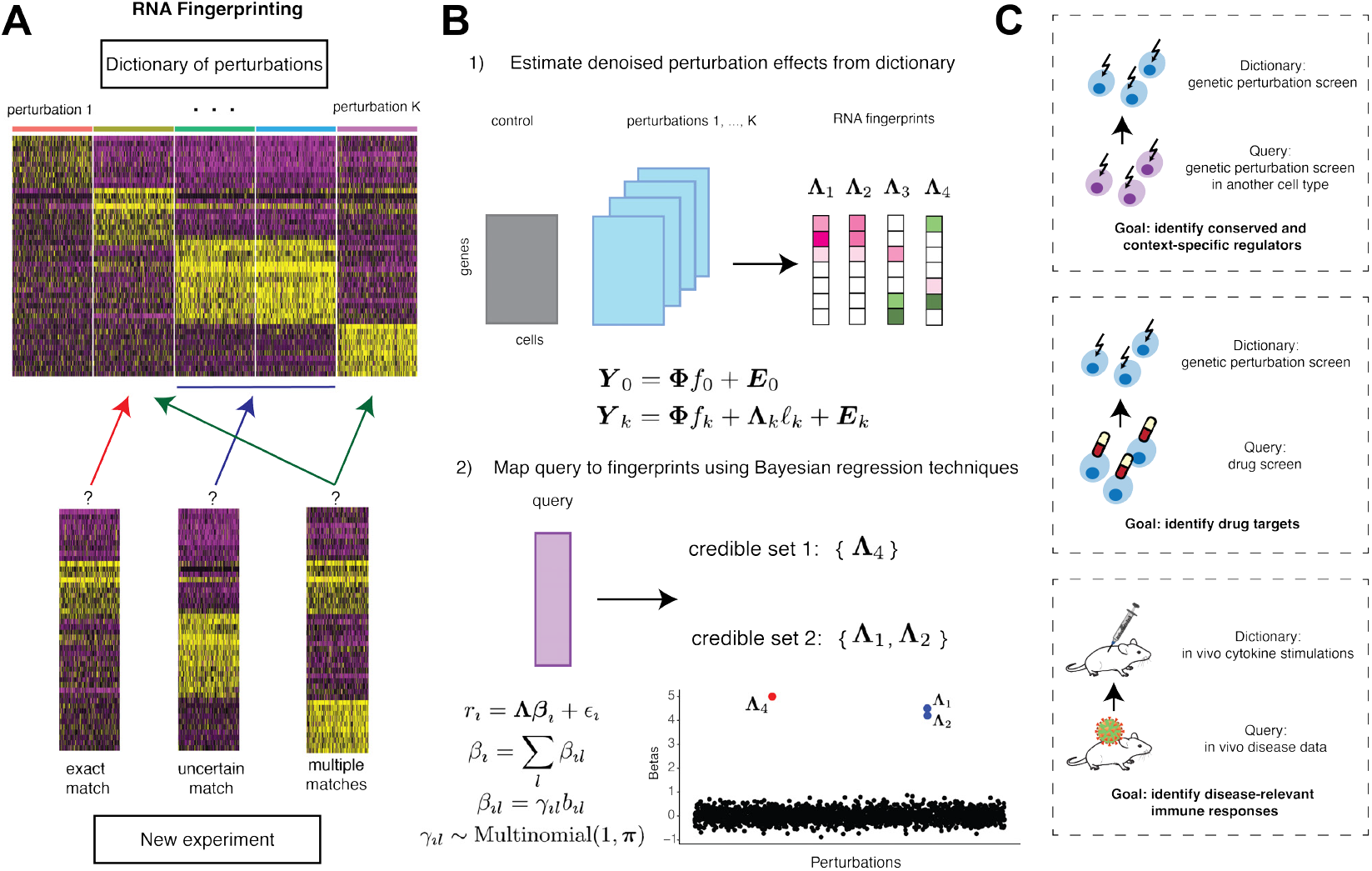
Overview of RNA fingerprinting. **(A)** Conceptual schematic of RNA fingerprinting. The method leverages an scRNA-seq dictionary of known perturbations (top), and (bottom) aims to match query data (where the true perturbation is unknown) to the transcriptional signatures of one or more perturbations. **(B)** The two-step statistical framework of RNA fingerprinting. First, denoised perturbation effects are estimated from the reference dataset to obtain a set of RNA “fingerprints” using a multi-condition latent factor model. Second, query data are mapped onto these fingerprints using Bayesian regression techniques adapted from statistical genetics, which yields credible sets of one or more fingerprints that best explain the query profile. **(C)** Example applications of RNA fingerprinting. Examples include comparing genetic perturbation screens across cell types to identify conserved and context-specific regulators, mapping drug perturbations to genetic perturbation fingerprints to identify drug targets, and mapping in-vivo disease datasets to cytokine stimulation dictionaries to identify disease-relevant immune responses.

An effective method for perturbation mapping, particularly at single-cell resolution, must overcome substantial statistical challenges. Both reference and query datasets can exhibit high levels of biological and technical variation, cell state heterogeneity, and variability in perturbation response strength^28,29^. These challenges are compounded when working with large dictionaries containing hundreds or thousands of perturbations, many of which may only have subtle transcriptional differences from one another, such as knockdowns of genes within the same complex. In such cases, a robust framework should allow for uncertainty in the absence of sufficient evidence for a definitive match. Finally, biological systems frequently exhibit mixtures of perturbation effects, requiring methods that can model combinatorial responses.

We address these desiderata through a two-stage statistical framework that we call RNA fingerprinting (Figure 1B; Methods). Briefly, the first stage infers denoised perturbation effect estimates, termed “fingerprints”, for each perturbation in the dictionary, then the second stage probabilistically assigns query cells to one or more of these fingerprints. In the first stage, we estimate a fingerprint for each perturbation using a multi-condition latent factor model (Methods). We account for confounding variation by estimating latent factors from control cells, which represent baseline sources of heterogeneity. For each perturbation, we then model the gene expression of the perturbed cells as arising from a combination of these baseline factors and a rank-one perturbation-specific factor, i.e. the fingerprint, that captures the primary transcriptional response to the perturbation and allows for the possibility of variation in response strength across cells. Each fingerprint can be interpreted as a denoised vector summarizing how the expression of each gene changes under the perturbation. The output of this stage is a dictionary of such fingerprints that have been disentangled from multiple forms of heterogeneity.

In the second stage of the framework, we map query data to one or more fingerprints that best explain their transcriptional responses. This mapping can be performed at the level of individual query cells, or by aggregating over groups of cells, such as those from a shared condition or cluster. After accounting for baseline variation through a similar procedure as above, we model each query’s gene expression as a linear combination of the learned fingerprints, where nonzero weights indicate contributions from specific perturbation effects. Estimating these weights can be framed as a regression problem, but a naive implementation, e.g. using ordinary least squares or ridge regression, would fail both due to the sparsity of relevant effects (where typically only a small subset of perturbations contribute meaningfully out of up to thousands considered), and because standard methods struggle to disentangle the effects of highly correlated features (such as fingerprints for regulators in the same complex).

To overcome these challenges, we draw an analogy to genome-wide association studies (GWAS), which face a similar problem of identifying causal single nucleotide polymorphisms (SNPs) among many highly correlated candidates. In GWAS, linkage disequilibrium often prevents pinpointing a single causal SNP^31–33^, which motivates reporting credible sets, i.e. groups of correlated variants that plausibly contain the true causal signal. Inspired by this approach, we adapt SuSiE^30^, a Bayesian variable selection method originally developed for GWAS finemapping, to report credible sets of fingerprints when a unique match cannot be confidently determined. Our method can return multiple such credible sets, each corresponding to a distinct transcriptional effect and containing one or more candidate fingerprints. Each set is accompanied by a Bayes factor quantifying the strength of evidence for its contribution. If no fingerprint in the dictionary sufficiently explains the query, the model returns no credible sets (unassigned), analogous to all regression coefficients being zero. This approach allows us to summarize complex and potentially ambiguous matches with interpretable probabilistic outputs.

Importantly, because SuSiE was built for GWAS data, there are some unique statistical challenges when adapting the model to single-cell RNA-sequencing (scRNA-seq). While SNP genotype data is discrete (dosage of 0, 1, or 2) and is measured with very low error, gene expression data is much noisier and has a far greater dynamic range. This gives rise to occasional extreme outliers that can substantially distort estimation. We address this by modifying the regression model to include additional terms that can adaptively absorb such extreme expression values (Methods). Moreover, our fingerprint estimates carry additional uncertainty due to the limited and variable number of cells available for each perturbation. We account for this in two ways: first, by explicitly estimating and propagating uncertainty from the fingerprint estimates when identifying credible sets, and second, by introducing a strategy to pool noisy perturbations together through proposals of joint effects (see Methods).

Our RNA fingerprinting framework yields three key outputs: first, denoised perturbation-specific signatures, or fingerprints, learned from the reference dictionary; second, one or more credible sets of fingerprints that best explain the transcriptional responses in each query cell or aggregated group; and third, a Bayes factor associated with each credible set, quantifying the strength of evidence for its contribution. Full implementation details are provided in the Methods, and we make our software publicly available at www.github.com/satijalab/rna-fingerprinting.

This framework enables flexible application across a wide range of biological questions (Figure 1C).

### RNA fingerprinting accurately maps perturbations within and across contexts

We first evaluated our framework in the context of ground-truth query datasets where the underlying genetic perturbation is known. As an initial demonstration, we analyzed the CPA-Perturb-seq dataset^34^, in which 42 regulators of cleavage and polyadenylation were knocked down using CRISPR interference (CRISPRi) across two distinct cell lines. This dataset is representative of common challenges for mapping, including heterogeneous perturbation efficiency and highly related perturbations targeting shared molecular complexes. We assessed our method by first estimating fingerprints from one replicate of the HEK293FT cells, and predicting the true perturbation identity from query cells in a second, independently performed replicate based on their expression profiles alone (Figure 2A).

**Figure 2.**
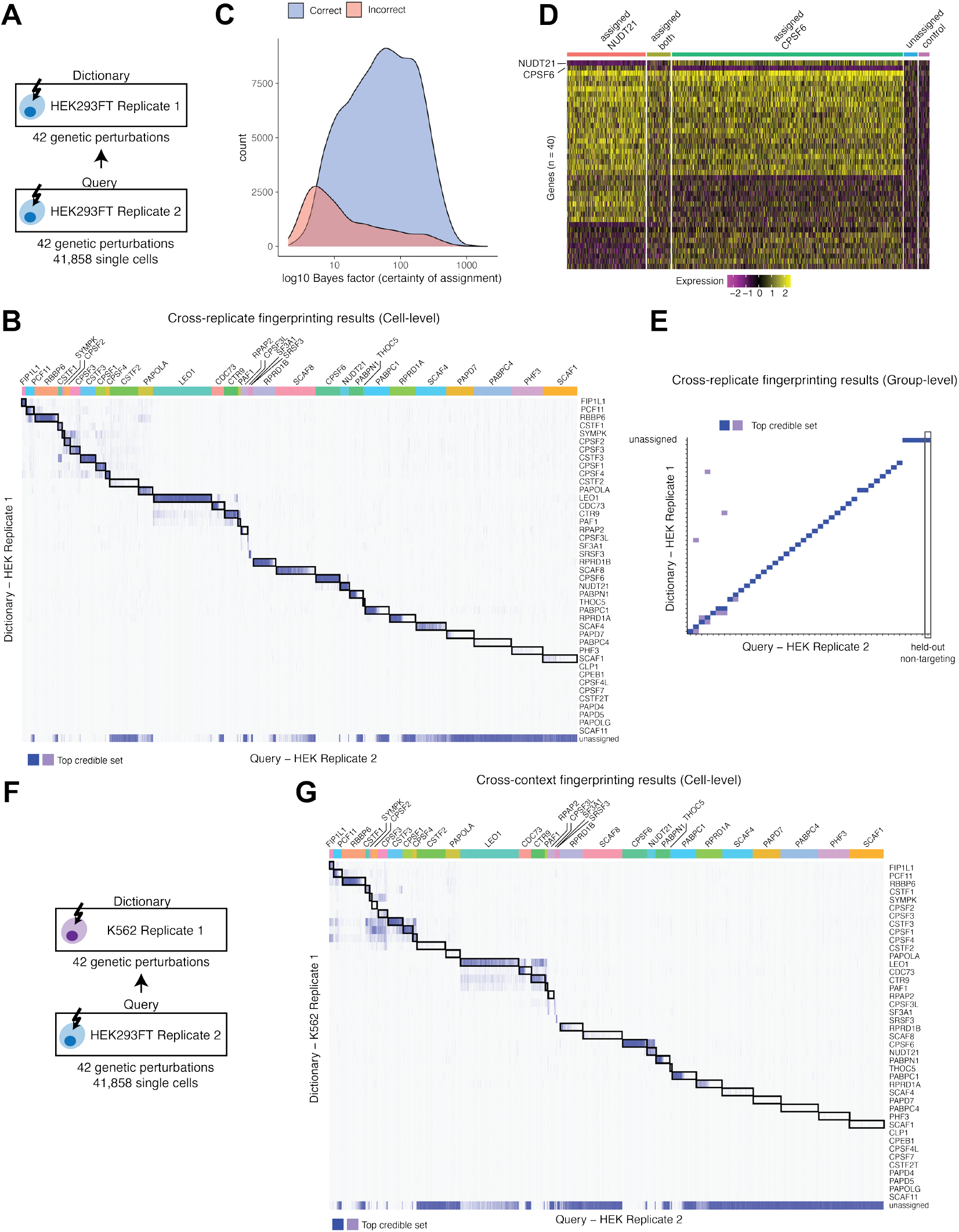
Demonstrating RNA fingerprinting within and across biological contexts in the CPA-Perturb-seq data. **(A)** Schematic of mapping across replicates in HEK cells from the CPA-Perturb-seq data, where data from Replicate 2 is mapped to fingerprints estimated from Replicate 1. **(B)** Heatmap of individual cell-level fingerprinting results across replicates. Each row corresponds to a reference fingerprint, and each column corresponds to a query cell, grouped by ground-truth labels. Shading denotes inclusion of the fingerprint in the query cell’s top credible set. Blue indicates the top credible set, and dark blue indicates the top member. Black boxes note the locations for ground-truth matches. **(C)** Distribution of Bayes factors (assignment certainty) for correct (blue) and incorrect (red) cross-replicate assignments, demonstrating strong separation between correct and incorrect matches. **(D)** Gene expression profiles (Pearson residuals) of cells receiving either NUDT21 or CPSF6 perturbations in the query data, grouped by whether they were assigned exactly “NUDT21,” exactly “CPSF6,” a credible set containing both labels, or no assignment at all. A random sample of non-targeting cells are also shown for comparison. **(E)** Group-level fingerprinting results for the top credible set across replicates. Heatmap structure and shading are the same as in (B). **(F)** Schematic of cross-context benchmarking, in which perturbations from a K562 replicate (dictionary) are used to assign cells from a HEK replicate (query). **(G)** Individual cell-level fingerprinting results across contexts, as in (B).

Examining the fingerprints learned from the reference replicate, we found that fingerprints clustered by complex as expected (Supplementary Figure 1A), but also that fingerprints learned independently for two separate guide RNAs (e.g. CPSF1, *R* = 0.90) exhibited clear reproducibility (Supplementary Figure 1B). Strikingly, fingerprint estimates were highly correlated even for subsets of genes that were not detected as differentially expressed. This demonstrates our ability to robustly quantify effects for subtly perturbed genes, with reproducible performance observed even when processing two guide RNAs for the same gene with differing knockdown efficacy (Supplementary Figure 1B). We observed reduced reproducibility (CPSF1, *R* = 0.79) when considering traditional bulk-based measures of perturbation strength (i.e. log fold-change), highlighting the value of explicitly modeling cellular heterogeneity, and our denoising approach more broadly.

Our learned fingerprints enabled us to accurately map perturbation identities for cells in the second replicate (Figure 2B). While we report results for all 42 perturbations in Supplementary Figure 1C, we focused on 33 perturbations that were found by the previous studies^28,34^ to introduce a transcriptomic change in at least some cells (Methods). RNA fingerprinting assigned perturbations to 15,918 cells out of 26,961 cells (59%). Assignments were accurate at single-cell resolution, with the correct perturbation being assigned in the top credible set in 88% of cases. Cells labeled as unassigned were consistent with expected heterogeneity in the effectiveness of perturbation in CRISPRi experiments^29^. For instance, while 35% of CDC73-perturbed cells were unassigned (Supplementary Figure 1D), they appeared to have escaped the perturbation as evidenced by a lack of expression knockdown of the target regulator as well as the broader transcriptomic profile (95% were labeled as “escaping” by Mixscape).

Assigned cells showed coherent transcriptional phenotypes with their labels (Supplementary Figure 2A). The correct assignments were highly specific (82% had a single credible set with only one fingerprint), and were associated with substantially increased Bayes factors compared to the small number of incorrect assignments (Figure 2C). In cases where larger credible sets were returned, they often corresponded to genuinely ambiguous transcriptional profiles (Figure 2D). For example, perturbations of NUDT21 and CPSF6, representing two of the most transcriptionally similar targets in the dataset, share hundreds of differentially expressed genes and differ only by subtle expression changes^34^. While the majority of cells perturbed with either gene were assigned the correct label with high confidence, a subset received a credible set containing both NUDT21 and CPSF6. Inspection of these cells suggests that their transcriptional profiles were consistent with either perturbation (Figure 2D). Taken together, these results demonstrate that RNA fingerprinting can return specific and accurate perturbation matches when supported by underlying data, but when faced with uncertainty, returns appropriate and interpretable credible sets.

We found that it was possible to further improve performance by pooling groups of single cells prior to classification (i.e. “group-level” matching). In this use-case, query cells continue to be processed individually by the latent variable model, but their responses are aggregated together based on relevant metadata, such as condition, class, or cluster, prior to running Bayesian regression. When aggregating query cells by their perturbation (Figure 2E), these “group-level” mappings were both highly accurate (95% correct out of assigned) and typically specific to a single perturbation (81% of correct assignments). These findings illustrate the power of group-level analysis to recover even very subtle perturbation phenotypes by borrowing strength across cells. We note that this application still requires single-cell data, as the query cells are individually corrected for baseline sources of heterogeneity prior to pooling – a key step that would not be possible when working with bulk RNA-seq data.

Finally, we evaluated the framework’s ability to map perturbations across distinct biological contexts by learning fingerprints from K562 cells and using them to annotate the same set of HEK293FT query cells (Figure 2F). These two cell types represent markedly different origins: K562 cells are derived from an erythroid leukemia, while HEK293FTs are epithelial-like cells derived from human embryonic kidney. This resulted in a higher unassigned rate at the singlecell level for the same set of perturbations (37% assigned). Nevertheless, RNA fingerprinting maintained high accuracy, with 81% of assigned cells including the correct perturbation in their top credible set (Figure 2G; full assignments in Supplementary Figure 2B). Notably, perturbations that failed to map reliably often exhibited divergent transcriptional responses between the two cell lines (Supplementary Figure 2C), suggesting that in some cases, the lack of correct assignment reflects a genuine biological limitation rather than a failure of the model.

Moving across contexts also increases the uncertainty of perturbation matches, represented by on average larger credible sets in assigned cells. For example, among cells receiving PAF-complex perturbations, 43% of assignments were both correct and specific (i.e. only one credible set returned, containing only the true member), while an additional 48% were mapped to credible sets containing one or more PAF-complex members (Figure 2G). This was also true at the group-level, where several results that had been specific within-context were now mapped to larger credible sets (Supplementary Figure 2D). RNA fingerprinting therefore may address the additional uncertainty in cross-context analysis by mapping at the level of molecular complexes when individual regulators cannot be confidently distinguished.

### RNA fingerprinting resolves combinatorial effects

While our previous analyses demonstrated the ability of RNA fingerprinting to identify a single causal perturbation, cellular responses are often shaped by multiple independent factors. We previously introduced CaRPool-seq, a Perturb-seq extension that induces and measures both single and dual-gene perturbations at single-cell resolution^35^. CaRPool-seq data of dually perturbed cells represents a ground-truth benchmark for assigning combinatorial effects: we mask the two known guide RNA identities to ask whether RNA fingerprinting can infer both perturbations from reference transcriptomic profiles of single perturbations (Figure 3A).

**Figure 3.**
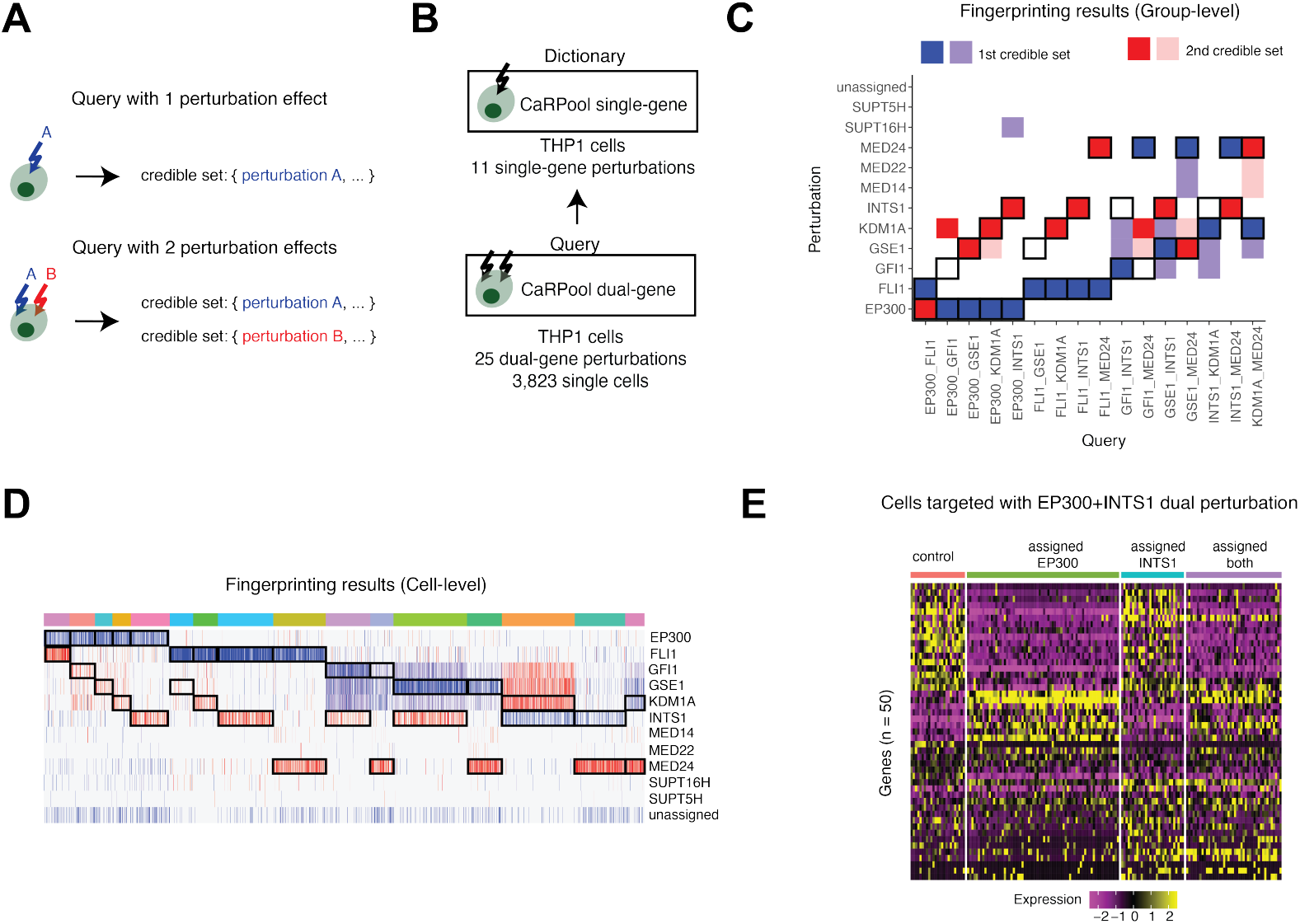
RNA fingerprinting identifies multiple responses in combinatorial screens. **(A)** Schematic of fingerprinting for single- versus dual-gene perturbations. A cell with one perturbation effect yields a single credible set, while a cell with two perturbation effects yields credible sets corresponding to both underlying perturbations. **(B)** Schematic of benchmarking using CaRPool Perturb-seq data in THP1 cells, where single-gene perturbations (dictionary) were used to assign cells with dual-gene perturbations (query). **(C)** Group-level fingerprinting results for dual perturbations. Heatmap shows the dictionary perturbations included in the first credible set (blue) or second credible set (red) for each query perturbation. Black boxes denote the expected ground-truth results. **(D)** Same as in (C), but showing fingerprinting results at the individual cell level. Blue shading reflects the first perturbation (ordered alphabetically) of the ground-truth label, and red shading reflects the second perturbation. Darker shade denotes the top member of the respective credible set. **(E)** Gene expression profiles (Pearson residuals) for cells targeted by the EP300+INTS1 dual perturbation, grouped by assignment to EP300, INTS1, both, or unassigned. Heatmap shows that the assignment from RNA fingerprinting reflects the cell’s underlying molecular phenotype.

To evaluate combinatorial perturbation matching, we used CaRPool-seq data from THP1 cells^35^ (Methods). We first constructed a reference dictionary of fingerprints for 11 single-gene perturbations, selected as those determined to have detectable transcriptional effects (Supplementary Figure 3A, Methods), then mapped the dual perturbations to the dictionary at both the group and single-cell levels (Figure 3B). Ideally, if the dual perturbations represent additive, independent effects, then mapping should return two credible sets, each containing one of the underlying perturbations.

When mapping all 25 profiled dual perturbations at the group level, RNA fingerprinting correctly matched to both perturbations in 80% of cases, but sometimes (32% of cases) mapped both in a single credible set (Supplementary Figure 3B). Focusing on the 16 dual perturbations in which the constituent effects were uncorrelated (Methods), we successfully found two credible sets, each corresponding to one of the underlying perturbations, in 11 of the pairs (69%, Figure 3C). In the remaining five cases, the model returned just one of the underlying perturbations: three times as a single credible set, and twice as two sets with only one correct match. Such cases can arise when one of the perturbations presents a stronger transcriptional effect, and the weaker signal is missed. These results demonstrate that RNA fingerprinting can successfully resolve combinatorial perturbations, but as expected, accuracy depends on the strength of the transcriptional signal.

RNA fingerprinting was able to map these combinatorial effects even at the single-cell level: 889 of 3,367 assigned cells (26%) returned two correct credible sets, while an additional 1,951 cells (58%) were assigned to just one of the two underlying perturbations (Figure 3D). Importantly, these assignments often reflected genuine heterogeneity rather than simple model error. For example, among cells where EP300 and INTS1 were simultaneously targeted, RNA fingerprinting assigned 37% to a single credible set containing EP300 and 23% to two sets respectively containing EP300 and INTS1. Visualizing single-cell expression profiles confirmed that only the latter subset exhibited a clear joint perturbation signature, whereas the former resembled the single-gene perturbation (Figure 3E). Similar patterns were observed for multiple dual perturbations (Supplementary Figure 3C). We conclude that RNA fingerprinting can identify combinatorial effects at single-cell resolution while distinguishing heterogeneity in perturbation outcomes.

### Genome-scale RNA fingerprinting

We next tested whether RNA fingerprinting could be extended to genome-wide scale by using Genome-Wide Perturb-seq (GWPS)^19^ as a reference. As one of the largest available perturbation dictionaries, this dataset profiled CRISPRi knockdowns of all 9,866 expressed genes in K562 cells, measured eight days after transduction. The same study also performed an independent experiment targeting 2,057 essential genes, collected at an earlier time point (six days), which we used as a query. We evaluated RNA fingerprinting’s ability to identify the correct perturbation in this essential-scale dataset, a challenging task given that there are thousands of possible candidates but only one true match in each case.

We learned a total of 4,098 fingerprints passing our filters (Methods) from the genome-scale experiment, then mapped cells from the essential-scale screen whose perturbations overlapped with these (Figure 4A). At the group-level, RNA fingerprinting gave assignments to 87% of query perturbations. Among these assigned perturbations, the top credible set contained the true label in 71% of cases, with accuracy increasing with increasing Bayes factor (Figure 4B). For our most confident calls, accuracy exceeded 90%. Moreover, the top credible sets remained specific even for this genome-scale dictionary, with a modal credible set length of 1 (i.e. RNA fingerprinting returned a single perturbation). These results highlight the sensitivity and precision of our approach, even in this large-scale context. One such example is the KAT8 perturbation, which displays a transcriptional fingerprint distinct from all other perturbations (Figure 4C). We can visualize this classification using “Long Island City” plots (Methods), inspired by GWAS Manhattan plots, which illustrate the specific identification of this perturbation.

**Figure 4.**
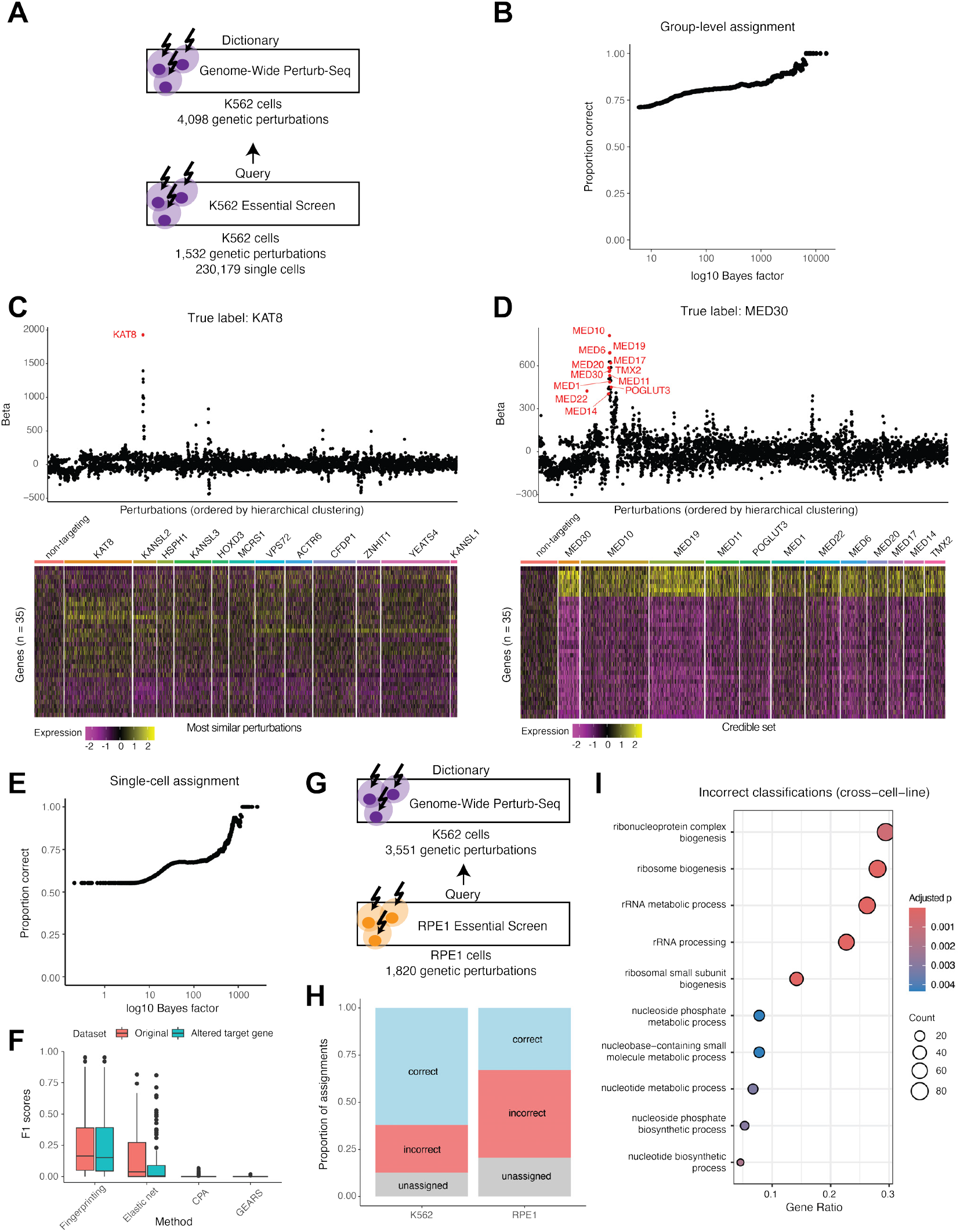
RNA fingerprinting at the genome scale. **(A)** Schematic of cross-dataset genome-scale benchmarking in K562 cells, where perturbations from a genome-wide Perturb-seq dataset were used as fingerprints for query cells from a separate screen of essential perturbations. **(B)** Group-level fingerprinting results, showing the proportion of correct assignments (defined as the right perturbation being present in the top credible set) out of all assignments with log10 Bayes factor greater than each x-axis value. **(C)** (Top) Long Island City plot (full description in Methods) for the KAT8-perturbed cells, mapped at the group-level. X-axis represents all possible perturbations grouped by hierarchical similarity, Y-axis reflects the posterior coefficient from Bayesian regression. Perturbations returned in credible sets are shown in red. (Bottom) Gene expression profiles (Pearson residuals) of KAT8-perturbed cells in the query, compared to non-targeting cells. Also shown are perturbations with the most similar profiles (i.e. the next highest conditional posterior mean coefficients), which were not assigned to a credible set. **(D)** As in (C), but for the MED30-perturbed cells. Additional perturbations shown are those in the top credible set. **(E)** As in (B), but for the individual cell-level fingerprinting results. **(F)** Boxplot of F1 scores for each perturbation in the benchmark subset of the query data, shown for RNA fingerprinting alongside comparator methods. For each method, scores are shown only for perturbations that were assigned at least once, as F1 scores cannot be calculated otherwise. For RNA fingerprinting and elastic net, we additionally report results after altering the target gene to ensure that its reduction via CRISPRi does not singularly drive the match (Methods). **(G)** Schematic of genome-scale mapping across cell lines. Same as in (A) but query experiment was performed in RPE1 cells. **(H)** Comparison of group-level assignment results in the RPE1 screen, versus the K562 screen. **(I)** Top enriched gene ontology terms for misclassifications in the RPE1 query data (Methods). Enriched terms for correct classifictions are shown in Supplementary Figure 4B.

While these results illustrate the potential for highly confident identification, not all perturbations are equally identifiable. A subset of essential-scale perturbations could not be assigned (13%), typically due to weak or undetectable transcriptional phenotypes in one or both datasets. In other cases, RNA fingerprinting returned long credible sets because multiple perturbations produced highly similar transcriptional profiles. This was particularly common for targets within large protein complexes, where redundancy in biological function translated into overlapping phenotypic signatures. For example, MED30, a mediator complex subunit, produced a longer credible set including nine other mediator members, reflecting their strongly correlated perturbation profiles (Figure 4D). Overall, our results show that RNA fingerprinting can resolve precise matches even from genome-wide reference dictionaries, though low signal strength and biological redundancy can still pose challenges for specific assignment.

We next mapped all 230,179 cells from the essential-scale screen individually. This represents an even more challenging scenario, given the prevalence of weak transcriptional phenotypes and perturbation escape. We assigned labels to 73,758 cells, of which 55% included the true label in the top credible set. As in the group-level setting, accuracy increased with Bayes factor (Figure 4E), indicating that higher-confidence calls were indeed more reliable. Most assignments were again highly specific, with a modal credible set length of one, and the perturbations yielding longer credible sets were similar to those observed previously. Notably, 96% of held-out non-targeting cells were correctly left unassigned, demonstrating strong false positive control even at this large scale. Thus, RNA fingerprinting can generate meaningful, robust single-cell assignments at genome-wide resolution.

To benchmark our performance, we compared RNA fingerprinting to three alternative singlecell assignment approaches: a penalized regression model (elastic net) and correlation-based matching to perturbation profiles generated by two deep learning methods, GEARS^36^ and CPA^37^ (Methods). While these other methods were not explicitly designed for this task, and therefore lack key features of RNA fingerprinting such as summarizing uncertainty via returning single or multiple credible sets, we sought to make the comparison as fair as possible. For each approach, we considered only the top predicted perturbation per cell; for RNA fingerprinting, this meant using the top perturbation from the top credible set even when more members were present.

Performance was summarized using perturbation-level F1 scores, which balance sensitivity and precision. RNA fingerprinting achieved the highest F1 scores, with elastic net ranking second (Figure 4F). Moreover, we found that elastic net assignments relied heavily on low target gene expression, an unreliable signal that is specific to CRISPRi perturbation dictionaries. When altering the query data to remove decreases in target gene expression (Methods), RNA fingerprinting showed only a slight drop in F1 scores, whereas elastic net performance fell sharply (Figure 4F, Supplementary Figure 4A). These results underscore the need for RNA fingerprinting: existing statistical methods fail to provide robust assignments, and current deep learning models, which are designed for predicting perturbation responses in new contexts, cannot be reliably repurposed for fingerprinting applications.

Finally, we tested cross-context performance by mapping an essential-scale screen from the same study performed in a different cell type, RPE1, to the Genome-Wide Perturb-seq fingerprints (Figure 4G). This application is particularly challenging given the substantial differences between the two lines, with RPE1 representing non-cancerous retinal pigment epithelial cells. We assigned labels to 1,444 of the 1,820 perturbations under consideration, of which 42% were correct (Figure 4H). Correctly assigned perturbations were enriched for core cellular processes (Supplementary Figure 4B), consistent with these responses being more likely to be conserved across divergent cell types.

We found that in many cases, when high certainty misclassifications occurred, they were made to a highly related perturbation. For example, some members of the proteasome complex were misassigned to credible sets containing other members of the same complex (Supplementary Figure 4C). Such patterns often arose when members of a complex exhibited weaker phenotypes in the K562 reference data but much stronger phenotypes in the RPE1 query set, leading the model to assign cells to other complex members with high Bayes factors to reflect the strong match. These results underscore the inherent challenges of cross-context classification, while also showing that even imperfect assignments can retain biological interpretability and suggest meaningful relationships.

### RNA fingerprinting identifies context-specific regulators of p53

To explore potential cell type–specific differences in perturbation function, we next considered cases where performance diverged sharply across contexts. We focused on perturbations that were correctly identified in the K562 essential-scale screen but were either unassigned or mapped to an unrelated perturbation in RPE1 cells. Enrichment analysis revealed that many of these targets were associated with ribosomal biogenesis, processing, and metabolism (Figure 4I). Within this group, RPL gene perturbations, which encode large ribosomal subunit components, stood out because they induced strong phenotypes in RPE1 cells yet matched their K562 counterparts in only a small subset of cases (Figure 5A).

**Figure 5.**
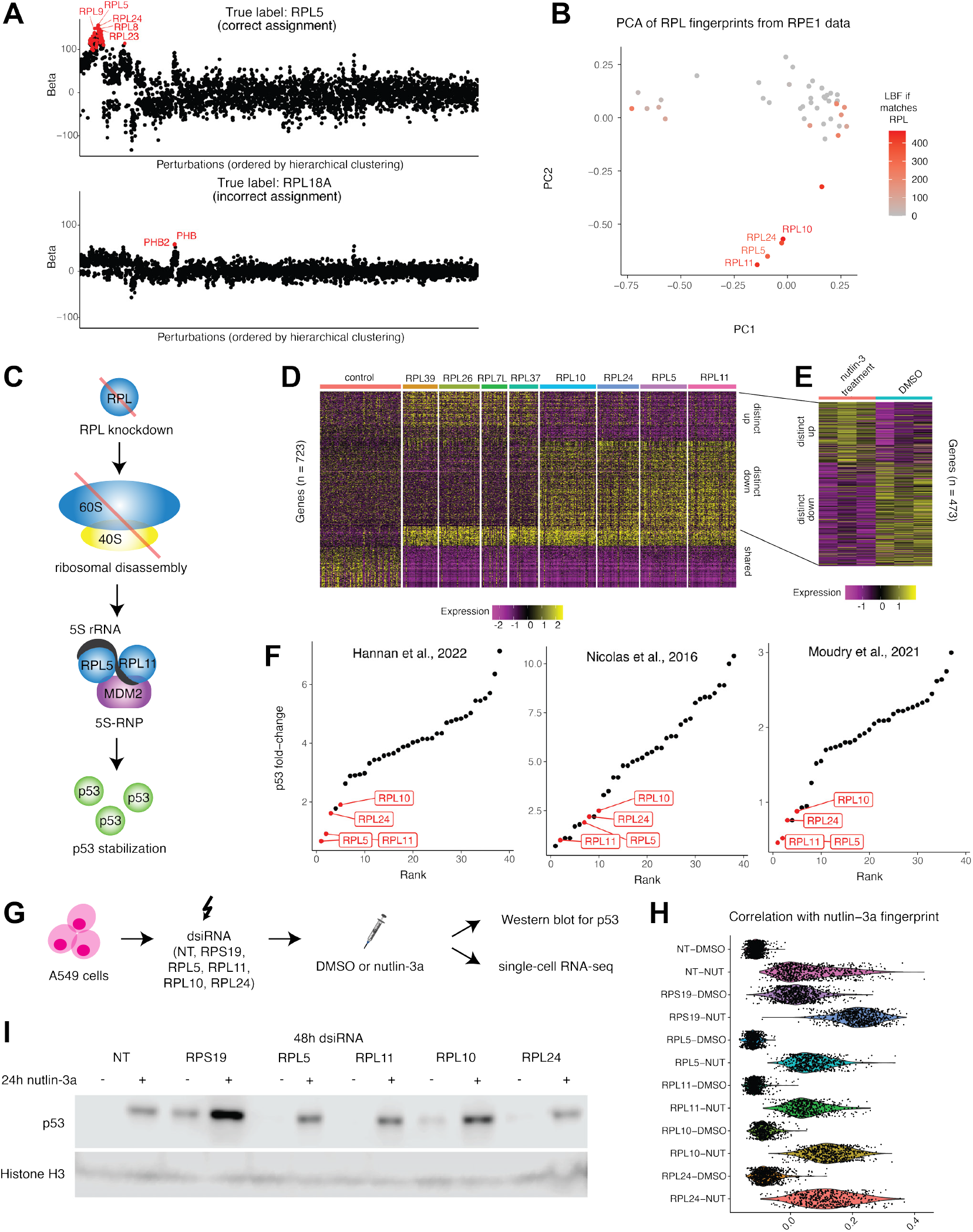
RNA fingerprinting links perturbation of RPL10 and RPL24 to p53 stabilization. **(A)** Long Island City plots of (top) RPL5-perturbed cells in the RPE1 query data, which successfully map to RPL fingerprints in K562 cells. (bottom) Same for RPL18A-perturbed cells, which do not map to RPL perturbations. **(B)** PCA plot of fingerprints corresponding to RPL perturbations estimated from the RPE1 data. Point color shows the log10 Bayes factor of perturbations who fingerprinted to an RPL perturbation in K562 cells, or are gray otherwise. **(C)** Schematic depicting canonical regulation of p53 via ribosomal stress. Ribosomal disassembly leads to formation of 5S-RNP (5s-rRNA + RPL5 + RPL11), which inhibits MDM2 and stabilizes p53. RNA fingerprinting results hypothesize that RPL10 and RPL24 are involved in this pathway. **(D)** Gene expression profiles (Pearson residuals) of RPE1 cells receiving knockdowns of selected RPL genes, showing shared and distinct transcriptional responses. Heatmap shows differential gene expression across successfully perturbed cells. **(E)** Comparison of distinct gene expression module in (D) to published signatures of nutlin-3a treatment vs. DMSO 45. **(F)** p53 protein levels after performing RPL perturbations that successfully induce ribosomal stress, as reported in three published datasets: Hannan et al. 40, Nicolas et al. 46, and Moudry et al. 47. **(G)** Experimental design for siRNA validation in A549 cells. Knockdowns of candidate RPL genes were combined with DMSO or nutlin-3a treatment, followed by Western blotting for p53 and single-cell RNA-seq. **(H)** Correlation of transcriptional profiles (Pearson residuals) between RPL knockdowns and the fingerprint estimated for nutlin-3a treatment (Methods). siRNA knockdown of RPS19 activates this module, but knockdown of RPL5, RPL11, RPL24, and RPL10 show defective activation. Additional treatment with nutlin-3a rescues this response. **(I)** Western blot validation in A549 cells. siRNA knockdowns of RPL5, RPL11, RPL24, and RPL10 in DMSO all lead to a failure to stabilize p53 protein, but RPS19 knockdown does not. MDM2 inhibition rescues p53 levels. Histone H3 serves as a loading control.

Fingerprint estimation in RPE1 cells revealed a distinct cluster of four perturbations (RPL5, RPL11, RPL10, and RPL24) that were matched with high confidence to RPL perturbations in K562 (Figure 5B). This was not the case for most of the remaining RPL perturbations, suggesting transcriptional divergence between K562 and RPE1 that may reflect cell type–dependent functional consequences. The presence of two groups provided an opportunity to understand why some RPL perturbations behaved so differently across cell types that they could not be matched, while others did not.

We reasoned that the differences between cell types may arise from their p53 mutational status: K562 cells are p53–null^38^, whereas RPE1 cells are p53–wild type^39^. Depletion of most RPL genes activates the ribosomal stress pathway, which induces a robust p53 response when ribosomes are impaired^40^. As a result, many RPL perturbations in RPE1 cells can trigger p53 activation, while the same perturbations in K562 cells cannot, creating systematic differences in their transcriptional profiles and making cross-context matching difficult. Notably, RPL5 and RPL11 are known to be essential for p53 activation in this pathway^41,42^. Under ribosomal stress conditions, RPL5 and RPL11 form the 5S ribonucleoprotein (RNP) particle, which binds MDM2 and prevents p53 degradation, in turn increasing p53 levels^43^ (Figure 5C). The depletion of RPL5 or RPL11 therefore blocks p53 induction even in p53–competent cells, potentially explaining their more concordant profiles between K562 and RPE1.

Our observation that RPL10 and RPL24 share highly similar fingerprints and are likewise matched with high certainty leads to a novel hypothesis for the function of these two genes. We propose that, in addition to their canonical role as members of the large ribosomal subunit, RPL10 and RPL24 may also be necessary for p53 activation in the presence of ribosomal stress. If correct, this would place them alongside RPL5 and RPL11 as key mediators of the p53 response in this pathway and results in a clear, testable prediction: perturbations of RPL10 or RPL24 should still induce ribosomal stress but fail to trigger downstream p53 activation (Figure 5C).

To test this prediction, we examined the transcriptional profiles of RPL perturbations in RPE1 cells. We identified a shared gene expression module present in nearly every RPL perturbation (Figure 5D) and linked this module to ribosomal stress by comparison with polysome profiling data from a previous perturbation screen^44^. Perturbations with high module expression showed reduced heavy polysome occupancy, as expected under ribosomal impairment, whereas those with low module expression displayed minimal changes (Supplementary Figure 5A). RPL10 and RPL24 perturbations expressed this module, consistent with the first element of our hypothesis that their depletion induces ribosomal stress.

Next, we asked whether perturbations of RPL10 and RPL24, like those of RPL5 and RPL11, fail to activate p53. We identified a gene module that distinguished these four RPL perturbations from others (Figure 5E). Strikingly, this module corresponded closely to previously published p53-responsive genes, as inferred from a bulk RNA-seq dataset of cells treated with the p53 activator nutlin-3^45^. Perturbations that induced ribosomal stress generally showed module expression consistent with p53 activation, while perturbation of RPL5, RPL10, RPL11, and RPL24 did not (Supplementary Figure 5B). Moreover, three independent screens measuring p53 protein abundance after RPL depletion^40,46,47^ consistently ranked these four among the lowest p53 inducers (Figure 5F). These results support the second element of our hypothesis, and explain how the misclassifications between K562 and RPE1 cells may stem from their different genetic capabilities.

Finally, to further validate and explore how RPL10 and RPL24 influence p53 levels, we performed dsiRNA-mediated depletion of each RPL in p53-competent A549 cells followed by scRNA-seq (Figure 5G). Cells were treated with either DMSO or the p53 activator nutlin-3a. Because nutlin-3a also functions by inhibiting MDM2, rescue of p53 activation by nutlin-3a would indicate a role for these RPLs in the 5S-RNP–MDM2 pathway, as opposed to a role independent of MDM2 inhibition. Indeed, after effective depletion of RPL10 (85% reduction) and RPL24 (91% reduction; Supplementary Figure 5C), activation of a module of p53-responsive genes was suppressed in DMSO but restored by nutlin-3a (Supplementary Figure 5D, Figure 5H). We performed western blots for p53 protein levels in each condition to confirm these results (Figure 5I). We observed similar results and rescue after depletion of RPL5 and RPL11, further supporting that all four regulators act within the same pathway. Finally, as expected, depletion of RPS19 as a positive control resulted in strong p53 induction even in the DMSO condition (Supplementary Figure 5D, Figure 5H-I). While additional work is needed to define the precise mechanism, our results strongly support that RPL10 and RPL24 are essential for 5S-RNP–mediated p53 activation under ribosomal stress. More broadly, these results demonstrate how RNA fingerprinting results can help discover context-specific differences in gene regulation.

### Mapping drug mechanism of action with RNA fingerprinting

Having demonstrated the utility of RNA fingerprinting for mapping genetic perturbations, we next asked whether the same framework could be applied to chemical perturbations. Prior work in the bulk context has compared sets of similar transcriptomic profiles between genetic and chemical perturbations to help characterize a drug’s mechanism of action^23,26^. We reasoned that RNA fingerprinting should be able to specifically map a drug treatment to the genetic perturbation of its true target. To test this, we aimed to match drug-treated cells from the sci-Plex multi-dose drug screen^20^ to genetic perturbations from Genome-Wide Perturb-Seq (CRISPRi knockdowns). Although these datasets were generated using distinct scRNA-seq technologies (sci-RNA-seq vs. 10x Genomics) and profiled at different timepoints (one day versus eight days), both experiments were performed in K562 cells, enabling direct comparison. We began with a set of three known BCR-ABL inhibitors (dasatinib, bosutinib, and nilotinib) that have been well-characterized. In each case, we mapped drug-treated cells at the group level to the reference dictionary of GWPS perturbations (Figure 6A).

**Figure 6.**
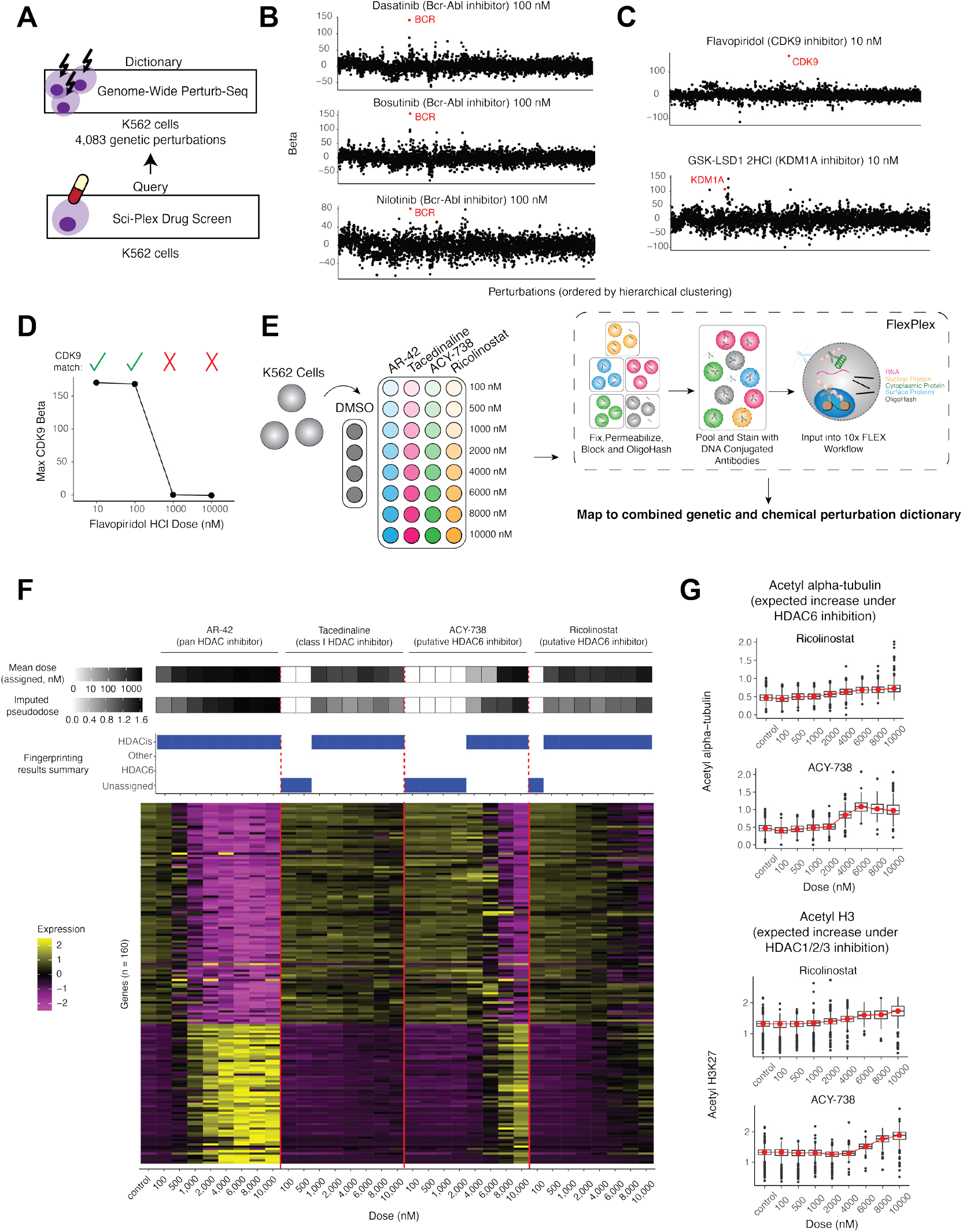
Mapping drug-treated cells to genetic perturbation screens. **(A)** Schematic of genetic (reference) vs. chemical (query) perturbation mapping. Reference fingerprints learned from Genome-Wide Perturb-Seq (GWPS) were used to map drug-treated perturbation profiles from sci-Plex. **(B)** Long Island City plots showing the top credible set from the group-level assignments of three Bcr-Abl inhibitor drug-dose combinations from the sci-Plex data. **(C)** Same as (B), but for two additional drug-dose combinations. **(D)** Dose–response fingerprinting of Flavopiridol. Mean CDK9 regression coefficient (as shown in the (C) y-axis), across doses demonstrates accurate mapping at low doses, with loss of specificity at higher doses. Checkmark denotes RNA fingerprinting assignment of CDK9 in the credible set. **(E)** Schematic illustrating the experimental design for HDAC inhibitor profiling. **(F)** (Top) Group-level classification results from the analysis illustrated in (E), summarizing the composition of each query’s top credible set (full results in Supplementary Figure 7B). For drug-dose combinations that mapped to HDAC inhibitors, we additionally show the mean dose of those fingerprints, as well as the imputed pseudodose (Methods). (Bottom) Pseudobulked gene expression for each drug-dose combination is shown for the genes in Supplementary Figure 8A. **(G)** FlexPlex intracellular protein measurements (collected alongside scRNA-seq) of acetyl alpha-tubulin and acetyl H3K27 for cells receiving each assessed dose of Ricolinostat or ACY-738.

Encouragingly, out of thousands of possible perturbations, RNA fingerprinting identified the correct genetic target, BCR, for each of the three drugs (Figure 6B). These assignments were highly specific: in each case, fingerprinting assigned a single credible set containing just the BCR CRISPRi perturbation in GWPS (Supplementary Figure 6A). Conventional approaches for comparing transcriptomic responses did not achieve this level of specificity. For example, two forms of gene set enrichment analysis on differentially expressed genes (Methods) identified a significant enrichment between BCR CRISPRi and dasatinib, but also returned significant results for hundreds of other unrelated perturbations. Similar non-specific matches were obtained using cosine similarity (Supplementary Figure 6B). Moreover, for all three of these approaches, BCR was not the top hit, in contrast to the correct and specific result returned by RNA fingerprinting.

We then considered the other drugs profiled in the sci-Plex screen. Many of these drugs are known to inhibit functional classes of proteins rather than a specific target, which may not match individual genetic knockdowns in GWPS. However, we identified two other drugs – a CDK9 inhibitor and a KDM1A inhibitor – where RNA fingerprinting again identified the correct target with complete specificity (Figure 6C). We confirmed that in both cases, gene set enrichment analysis and cosine similarity were unable to return a specific match (Supplementary Figure 6C,D).

Despite these overall successes, there were doses among these drugs at which the correct target was not recovered, reflecting underlying dose-dependent changes in transcriptional response. For example, RNA fingerprinting correctly identified CDK9 as the target of Flavopiridol at the two lowest doses, but not at the two highest doses. Gene expression analysis across doses revealed a marked shift in transcriptional profile at the higher doses (Figure 6D, Supplementary Figure 7A), consistent with altered specificity, and demonstrating that RNA fingerprinting can capture dose-dependent shifts in drug selectivity.

We next applied RNA fingerprinting to investigate potential off-target effects of HDAC6 inhibitors. Broad-spectrum HDAC inhibitors (HDACis) simultaneously target multiple classes of histone deacetylases (HDACs) for treatment of a wide variety of cancers, but more recent efforts have focused on more selective inhibitors to reduce toxicity^48,49^. Although HDAC6 inhibitors are intended to act specifically on the HDAC6 enzyme, previous studies suggest that some may lack true selectivity^50,51^. HDAC selectivity is typically evaluated by measuring levels of canonical histone acetylated substrates, but we reasoned that RNA fingerprinting could provide a broad, complementary, and rich basis for assessment.

We focused on two putative HDAC6 inhibitors, ACY-738 and Ricolinostat^52^. We individually treated K562 cells with these drugs as well as with two positive controls: the pan-HDAC inhibitor AR-42 and the class I HDAC inhibitor Tacedinaline, which primarily targets HDAC1, HDAC2, and HDAC3. All drugs were profiled across a range of eight doses followed by scRNA-seq using FlexPlex, our recently developed workflow^53^ (Methods), to capture both transcriptomes and canonical intracellular protein markers of HDAC inhibition (Figure 6E). Specifically, we measured acetylated alpha-tubulin as a marker of HDAC6 inhibition, and acetylated histones as a marker of class I HDAC inhibition^54^. These protein measurements provide an orthogonal benchmark for our RNA fingerprinting assignments and a direct comparison to the conventional selectivity assessments based on protein markers.

We then mapped these drug-treated cells to a combined dictionary containing both genetic and chemical perturbations (Figure 6E). This dictionary included individual CRISPRi knockdown fingerprints for 10 individual HDAC perturbations, including data from GWPS supplanted with additional Perturb-seq profiles from this study (Methods). We also included fingerprints from 311 drug–dose combinations from the sci-Plex screen, 64 of which corresponded to HDAC inhibitors, and therefore represented a proxy for non-additive combinatorial interactions for which Perturbseq data was not available. If Ricolinostat and ACY-738 act as true HDAC6-selective inhibitors, they should match to fingerprints of HDAC6 inhibition; if instead they are non-selective, we would expect assignments to fingerprints corresponding to other broad-spectrum HDAC inhibitors in the dictionary.

When first considering our positive controls AR-42 and Tacedinaline, RNA fingerprinting successfully matched these to HDAC inhibitor fingerprints in the dictionary, as opposed to alternative drugs or individual HDAC genetic perturbations (Supplementary Figure 7B). We successfully made these matches even at doses with subtle transcriptional phenotypes, such as the lowest dose of AR-42 (Figure 6F). Higher doses of query drugs tended to match fingerprints of higher doses among the reference drug fingerprints as well. This association was present for both measured dose but also “pseudodose”, defined by the original sci-Plex manuscript^20^ as the progression along a transcriptionally-defined continuum of broad HDAC inhibition (Supplementary Figure 7C). We imputed pseudodose values for each query drug based on its RNA fingerprinting matches (Figure 6F). We found that the highest doses of AR-42 mapped to the highest pseudodose levels, while Tacedinaline treatment saturated at intermediate pseudodose levels, consistent with its more mild transcriptional phenotype.

We next considered the putative HDAC6 inhibitors Ricolinostat and ACY-738. Neither drug matched to HDAC6 knockdown fingerprints at any of the eight tested doses (Supplementary Figure 7B, Figure 6F). We note that our dictionary included Perturb-seq data from 4 different independent gRNA, as well as GWPS data, to mitigate the possibility of ineffective or off-target gRNA precluding a match. Instead, RNA fingerprinting classified these drugs as either unassigned (at low dose) or matched them to broad non-selective HDAC inhibitor fingerprints (Supplementary Figure 7B, Figure 6F). As we observed with the positive controls, increased drug concentrations were generally associated with higher imputed pseudodose, driven by increased transcriptomic progression along the previously defined trajectory. Consistent with this, transcriptomic signatures associated with this trajectory did not overlap with CRISPRi perturbation of HDAC6, but partially overlapped with bulk published RNA-seq data of HDAC1/2/3 triple knockout in a different cell line^55^ (Supplementary Figure 8A). Taken together, these results suggest that, under the conditions tested in this biological context, these compounds do not produce a transcriptional phenotype consistent with selective HDAC6 inhibition.

The protein marker measurements from FlexPlex further supported our findings (Supplementary Figure 8B). As expected for our positive controls, AR-42 treatment led to an increase in both acetylated histones and acetylated alpha-tubulin, while Tacedinaline increased only acetylated histones. Consistent with RNA fingerprinting, Ricolinostat and ACY-738 generally increased both markers across doses, as would be expected under broad HDAC inhibition rather than selective HDAC6 targeting (Figure 6G). Notably, at 4,000 nM, ACY-738 markedly increased alpha-tubulin acetylation without changing histone acetylation, a pattern that would conventionally suggest HDAC6 selectivity, yet RNA fingerprinting still matched this profile to low-dose broad-spectrum HDAC inhibitors (Supplementary Figure 7B). While we cannot definitively rule out that this drug exhibits HDAC6 selectivity at this dose, the totality of transcriptomic evidence suggests a pattern of broader HDAC inhibition. We conclude that RNA fingerprinting can be used to model chemical and genetic perturbations and can make predictions regarding drug sensitivity, specificity, and dosage that are validated by orthogonal evidence.

### Quantifying immune response in influenza-rechallenged mice

While our previous analyses focused on in-vitro perturbation dictionaries, we next aimed to demonstrate RNA fingerprinting in-vivo. These applications benefit from the single-cell resolution of RNA fingerprinting, as cells within living organisms may mount heterogeneous responses not only across cell types, but even within a single cell type. Our strategy is enabled by recently generated in-vivo perturbation datasets such as the Immune Dictionary^22^, which profiled the responses of 18 cell types to 88 cytokines in the mouse lymph node. We reasoned that RNA fingerprinting could use this dictionary to help identify the causal cytokines that drive heterogeneous immune responses in-vivo.

As a case study, we re-analyzed data from MacLean et al. ^56^, which investigated how the murine lung mounts rapid immune responses during secondary influenza infection. The authors showed that upon viral rechallenge, a subset of resident memory B cells migrate to infection sites to initiate local antibody production. Moreover, depletion of alveolar macrophages with clodronate-loaded liposomes (CLLs) abrogated the response, highlighting the role of intercellular communication networks. The study also generated scRNA-seq profiles of lung immune cells under three conditions: uninfected control (UI), secondary infection (SI), and secondary infection with macrophage depletion (SI+CLL), which we used as query datasets.

We applied RNA fingerprinting by first learning cytokine response fingerprints for each cell type from the Immune Dictionary (Methods), then mapping murine lung cells from the SI and SI+CLL conditions to this dictionary at single-cell resolution (Figure 7A). Of 8,498 query cells, 6,670 (78%) were assigned a cytokine response, with the remainder left unassigned. Response rates varied across both cell types and conditions (Figure 7B). Monocytes/macrophages (98%) and neutrophils (92%) showed the highest rates in both query states, whereas B cells exhibited a clear shift: 67% of cells mapped to a cytokine response in SI mice, but only 27% in SI+CLL. This suggests that depletion of alveolar macrophages, which is known to block B cell migration, also specifically disrupts the cytokine-driven responses of this population.

**Figure 7.**
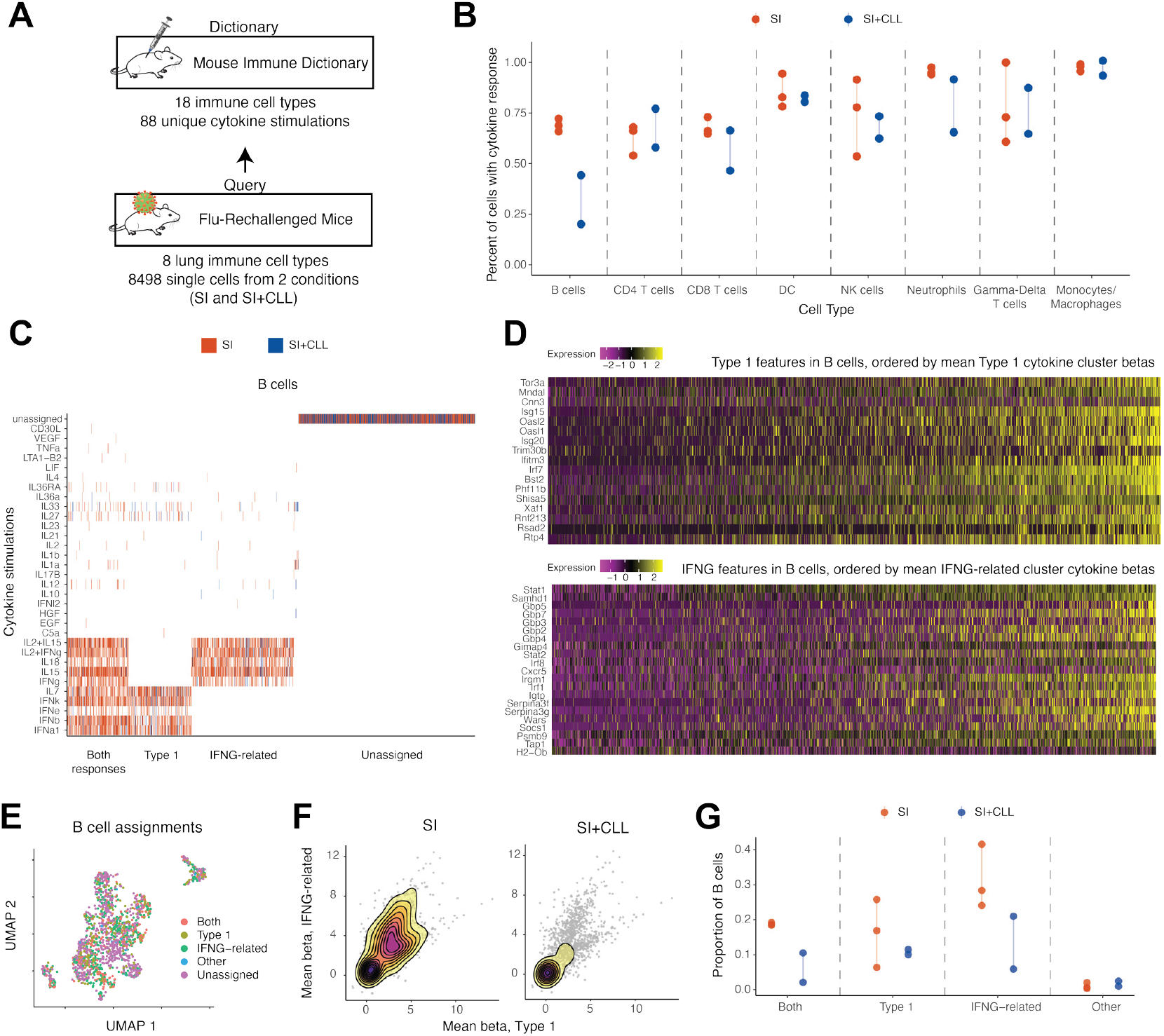
Quantifying immune response in influenza-rechallenged murine lung cells. **(A)** Schematic of in-vivo RNA fingerprinting. Reference cytokine perturbation profiles from the Immune Dictionary were used for mapping influenza-rechallenged immune cells from the mouse lung. **(B)** The proportion of cells exhibiting a cytokine response, shown for each cell type and mouse, and split by condition (SI or SI+CLL). Y-axis shows the proportion of cells mapping to one or more cytokine stimulations in the Immune Dictionary. Each dot represents data from one mouse, and the vertical lines denote the range of values. A small amount of random vertical jitter is added to separate overlapping points. **(C)** RNA fingerprinting results showing members of the top credible set for each query B cell. Individual cells are represented as columns (colored by condition), and are clustered into groups based on assignments. **(D)** Gene expression profiles (scaled expression) of all SI and SI+CLL B cells, showing selected genes representative of assignment to (top) the Type 1-IFN cytokine cluster and (bottom) the IFNG-related cytokine cluster. Cells are ordered separately for each heatmap, based on their average cytokine response (Methods). **(E)** UMAP visualization of unsupervised analysis of B cells from SI and SI+CLL mice, colored by cluster responses as defined in (C). **(F)** Strength of cytokine response programs in individual B cells. Each dot is a single cell, with x and y-axis values corresponding to regression coefficients for IFNG-related and Type 1 IFN fingerprints. Overlaid densities depict shifts between cells from SI (left) or SI+CLL (right). **(G)** The proportion of B cells belonging to each assignment cluster in each mouse, split by condition.

We next aimed to identify specific cytokines that may be involved in driving differential B cell responses between the SI and SI+CLL conditions. The assignments revealed heterogeneity that could group nearly all credible sets (98%) into three main cytokine response states (Figure 7C). One cluster mapped to primarily Type I interferons (e.g., IFNA1, IFNB), another to IFNG-related signals (including IFNG, IL-15, and IL-18), and a third showed a mix of both types of responses. Combined with the set of cells marked as unassigned, these results suggest that invivo cytokine responses within the murine lung are complex and non-uniform. Examining genes associated with the IFNA1/IFNB and IFNG fingerprints (Supplementary Figure 9A) confirmed this: both types of cytokine response programs represented molecular axes of variation within the B cells (Figure 7D). When applying the immune response enrichment analysis^22^ provided by the Immune Dictionary to B cells (Methods), the top cytokines were concordant with our results, but 69 total cytokines were returned as significantly enriched (FDR *<* 0.05).

Importantly, the identification of heterogeneous cytokine responses could not be obtained under existing approaches. Enrichment analysis strategies (which are based on differential expression of previously defined binary groups) cannot be used to identify new sources of heterogeneity within a cell type. Moreover, B cell subsets were not detected in standard unsupervised scRNA-seq analysis (Figure 7E). Even after subclustering B cells, unsupervised analyses primarily reflected other sources of variation (Supplementary Figure 9B), including differentiation state and kappa versus lambda chain usage, none of which separated our identified B cell subsets (Supplementary Figure 9C).

We further identified that these B cell subsets were linked to alveolar macrophage depletion. We used the fingerprinting model outputs to quantify each response type (Type 1 or IFNG) in the B cells (Methods) and observed a clear reduction of cytokine activity, particularly along the IFNG axis, upon the depletion of alveolar macrophages (SI+CLL mice) compared to SI mice (Figure 7F). We then compared the proportions of B cells assigned to each cluster of response across mice (Figure 7G). Although sample size was limited, differences between SI and SI+CLL mice were apparent only in the IFNG-related and mixed-response populations, but not in the primarily Type I interferon population.

Taken together, these results highlight a central role for IFNG in driving B cell responses during secondary infection. Although derived from computational predictions, they are consistent with the findings of MacLean et al. ^56^, who showed that neutralizing IFNG prior to secondary infection sharply reduced B cell migration. Whereas their analysis focused on differential expression of the Ifng transcript, RNA fingerprinting detects the downstream effects of cytokine signaling. By leveraging transcriptome-wide responses rather than a single gene, our complementary approach can increase robustness to low transcript abundance or RNA–protein discrepancies, both of which are common in immune signaling. While additional work, including spatial analyses, will be needed to determine whether the IFNG-responding B cell population corresponds to the migrating subset observed in vivo, we conclude that RNA fingerprinting successfully pinpointed contributing cytokines and revealed heterogeneous in vivo responses not detectable with conventional workflows.

## DISCUSSION

We introduce RNA fingerprinting, a statistical framework for mapping transcriptional responses from new single-cell experiments onto large-scale perturbation dictionaries. By learning denoised representations of perturbation effects and probabilistically assigning query cells to credible sets of reference perturbations, our approach enables causally-motivated interpretation of cellular responses across diverse experimental settings. We demonstrate its accuracy on ground-truth datasets and scalability to genome-wide screens. We additionally show that RNA fingerprinting can resolve additive combinatorial effects, identify drug mechanisms of action and dose-dependent selectivity, and characterize heterogeneous cytokine-driven immune responses in vivo. Together, these results establish RNA fingerprinting as a general framework for interpreting new single-cell data through the lens of perturbation reference dictionaries.

While the problem of matching transcriptional signatures has been widely explored in bulk contexts^23–25^, our single-cell approach offers unique advantages. Learning perturbation fingerprints directly from single-cell data enables us to explicitly control for confounding sources of variation, such as heterogeneous perturbation efficacy or cell state differences, that cannot be resolved with bulk RNA-seq. As a result, our framework yields precise and reproducible fingerprints that capture subtle perturbation effects more robustly than conventional measures, while also allowing nuisance variation to be subtracted from query cells before mapping at either the group or single-cell level. We believe that single-cell resolution for both reference and query datasets is essential to the success of this approach – and its rapidly growing availability across diverse perturbation contexts makes this strategy increasingly feasible.

A second key advantage of RNA fingerprinting is its probabilistic assignment framework. Rather than forcing each cell to match a single perturbation, the model returns credible sets of candidate fingerprints, together with Bayes factors that quantify the strength of evidence. We demonstrate the importance of this capability across a range of applications, and note that it arises from our Bayesian linear modeling framework. Utilizing nonlinear modeling strategies or deep neural networks for RNA fingerprinting would represent exciting extensions for future work, though doing so may sacrifice model interpretability and uncertainty estimation. Our work introduces multiple quantitative benchmarks with known ground truth, establishing a foundation for the systematic evaluation of future methods.

The availability of appropriate and context-matched perturbation dictionaries is a current limitation for broad in-vivo analysis, but the improving cost and scale of scRNA-seq continues to facilitate rapid data generation both in-vitro and in-vivo. In addition to genome-wide resources, our work highlights the pressing need for combinatorial perturbation dictionaries. While RNA fingerprinting can map additive signals, it is not designed to map synergistic responses. However, even the most specialized models struggle to robustly predict combinatorial synergy^57–60^, necessitating the generation of combinatorial perturbation datasets. Emerging technologies such as CaRPool-seq^35^, CROP-seq-multi^61^, and CRISPRAI^62^ now make it feasible to generate such resources at scale. Our analyses suggest that these efforts will be transformative, particularly for applications like determining drug mechanism of action, where compounds often act through multiple interacting targets rather than a single gene.

Finally, a fundamental assumption of our approach is that cellular transcriptomes encode perturbation responses, but RNA is only one limited aspect of cellular state. We propose this as both a current limitation and a promising opportunity for future work to extend to additional modalities. These include chromatin accessibility, post-transcriptional processing such as alternative polyadenylation and splicing, and protein abundance, all of which could be incorporated within the same modular framework by tailoring modality-specific noise models to each data type^34^. Incorporating morphological features from imaging datasets represents a particularly compelling opportunity, as this would enable the analysis of emerging and massively scalable optical pooled screening datasets^63^. Ultimately, we envision single-cell fingerprinting as the foundation for analyzing a new generation of multimodal reference maps that will transform how perturbation responses are interpreted across biology.

## DATA AND CODE AVAILABILITY

Previously published datasets were accessed as described in the Methods. Data generated in this study are made available at.https://doi.org/10.5281/zenodo.17127744. RNA fingerprinting is made available as an open-source R package at.www.github.com/satijalab/rnafingerprinting.

## ACKNOWLEDGMENTS

We thank all the members of the Satija Lab and Knowles Lab for thoughtful discussions related to this work. We would also like to thank Dr. Timothy McKinsey, Dr. Denis LaFontaine, Dr. Tal Arnon, and Samuel Woolliscroft for helpful discussions related to the biological applications, and Dr. Ang Cui for providing metadata for the Immune Dictionary data. I.N.G. is the Kenneth G. Langone Quantitative Biology Fellow of the Damon Runyon foundation (DRQ-21-24). J.D.B. is a postdoctoral fellow of the Jane Coffin Childs Memorial Fund for Medical Research. This work was supported by the Chan Zuckerberg Initiative (EOSS-0000000082, HCA-A-1704-01895 to R.S.), the NIH (RM1HG011014-02, 1OT2OD026673-01, R01HD096770, R35NS097404 to R.S), and the MacMillan Family and the MacMillan Center for the Study of the Non-Coding Cancer Genome at the New York Genome Center.

## AUTHOR CONTRIBUTIONS

I.N.G., D.A.K., and R.S. conceived of the study and wrote the manuscript. All experiments were performed by J.L., J.D.B, C.D., I.M., and A.B. I.N.G. performed computational analysis with supervision from D.A.K. and R.S.

## DECLARATION OF INTERESTS

In the past 3 years, R.S. has received compensation from Bristol Myers Squibb, ImmunAI, Resolve Biosciences, Nanostring, 10x Genomics, Parse Biosciences and Neptune Bio. R.S. is a co-founder and equity holder of Neptune Bio.

## METHODS

### RNA fingerprinting statistical framework

Our framework takes two datasets as input: a reference perturbation dictionary and a query dataset. The perturbation dictionary should consist of gene expression data for cells that either received no perturbation or one of a set of perturbations. The query dataset should also include a set of control or baseline cells. Our framework then maps the remaining cells in the query data to perturbations in the dictionary, returning assignments either at the individual cell level or at the group level by aggregating cells according to a label of interest (e.g. condition, treatment, cluster, etc.). RNA fingerprinting has two main stages. In the first stage, we estimate “RNA fingerprints” for each perturbation in the reference dictionary using a multi-condition latent factor model. In the second stage, we then map each query cell or group by fitting a Bayesian regression model in terms of these fingerprints. The methods are implemented in an open source R package available at.www.github.com/satijalab/rna-fingerprinting. Below, we describe these steps in detail.

#### Learning fingerprints with a multi-condition latent factor model

We begin by estimating fingerprints, or denoised representations of perturbation effects, for every reference perturbation in the dictionary. This is done by fitting a multi-condition latent factor model, applying an iterative outlier correction procedure, quantifying uncertainty in each set of estimates, and optionally filtering out low-signal fingerprints. These steps, detailed below, are implemented in the function LearnDictionary in the R package.

Suppose we have gene expression data from a perturbation dictionary consisting of *G* genes and *N* cells, where every cell either received a control perturbation (e.g. non-targeting guides or DMSO) or one of *K* possible perturbations. We first normalize this data using SCTransform v2^64,65^ with default parameters as implemented in the R package Seurat^66^, which applies a negative binomial modeling framework for normalization and variance stabilization. We then select a subset of *G*\* genes to use for fingerprint estimation; for example, this could consist of the union of top differentially expressed genes for each perturbation, or could be the set of all genes with average expression above some threshold (implemented as ChooseFeatures in the package). We describe the exact choices that we used for each analysis in this work in the corresponding sections below.

Let ***Y*** represent the resulting *G*\* *× N* matrix of SCTransform residuals, *S*_0_ be the set of column indices of the *N*_0_ control cells, and *S*_*k*_ those of the *N*_*k*_ cells that received perturbation *k*. Then

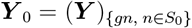

represents the columns corresponding to the control cells,

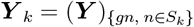

the columns for peturbation *k*, and we assume the multi-condition latent factor model,

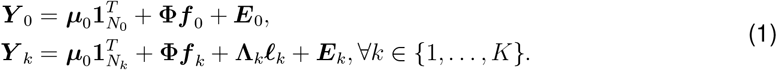

This model accounts for two primary sources of heterogeneity in the perturbed cells: baseline heterogeneity, which occurs in both perturbed and control cells, and variation in the strength of the perturbation, which is specific to the perturbed cells. We assume *M* baseline latent factors, so **Φ** is a *G*\* *× M* matrix of latent factor loadings encoding sources of baseline variation (e.g. cell cycle, differentiation stage, and other aspects of baseline heterogeneity), with corresponding *M × N*_0_ factor scores ***f*** _0_ for the control cells and *M × N*_*k*_ scores ***f*** _*k*_ for the cells that received perturbation *k*. Each entry of ***f***_**0**_, ***f***_***k***_ represents the contribution of each shared factor to each cell. The fingerprint **Λ**_*k*_ is a *G*\* *×* 1 latent factor loading encoding the effect of perturbation *k*, with the 1 *× N*_*k*_ score vector ***ℓ***_*k*_ denoting the strength of contribution to each cell. Finally, ***µ***_0_ is a *G*\* *×* 1 vector denoting baseline mean expression of each gene, and *G*\* *× N*_0_ matrix ***E***_0_ and *G*\* *× N*_*k*_ matrix ***E***_*k*_ represent random noise.

Under this model, the perturbation effects **Λ**_*k*_ are disentangled from baseline heterogeneity, and linear variation in perturbation response strength is accounted for via the cell-specific scores ***ℓ***_*k*_. We refer to each **Λ**_*k*_ as the *fingerprint* for perturbation *k*, and can interpret its entries as describing the denoised expression change in each gene under the perturbation.

#### Parameter estimation

To fit the model in Equation (1), we first estimate ***µ***_0_ and **Φ** using only the control data ***Y*** _0_. We set the estimator 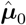 to the vector of gene-wise means of ***Y*** _0_. We then take the centered data 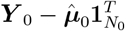 and apply principal components analysis (PCA) to estimate 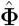 as the principal components and correspondingly 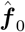 as the principal component scores. We use the truncated PCA implementation in the irlba R package^67^ for computational efficiency, and set the number of latent dimensions *M* = 30 by default.

Next, for each perturbation *k*, we estimate ***f*** _*k*_ by projecting the data ***Y*** _*k*_ onto the principal components 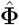. By default, we only consider perturbations with at least 10 cells. Specifically, we set

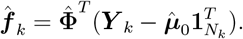

Then we can define the residual matrix

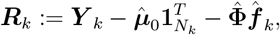

which represents the remaining expression in the perturbed cells that cannot be explained by baseline factors. We subsequently apply a rank-one PCA to the uncentered ***R***_*k*_ to estimate 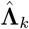 and 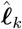 as the principal component and score vector respectively.

Because PCA is sign-invariant, the sign of the fingerprint 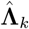 is arbitrary, i.e. we could equivalently estimate 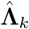 or 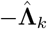 under this procedure. For downstream interpretability, we correct the sign of 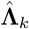 such that positive entries imply an increase in expression under perturbation *k*. This is done by first generating an analogous residual matrix of non-perturbed cells as

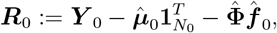

then applying the Wilcoxon rank-sum test to assess differential expression for each gene between ***R***_0_ and ***R***_*k*_, using the fast implementation provided in the presto R package^68^. If there are more than 1,000 non-perturbed cells, we randomly subsample the columns of ***R***_0_ to 1,000 cells for computational efficiency. We then compute the correlation between 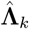 and the resulting vector of Wilcoxon rank-sum test statistics, and multiply 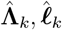 by *−*1 if this correlation is positive.

#### Iterative outlier correction

Our model assumes that the residual expression ***R***_*k*_ can be entirely attributed to either the rank-one perturbation effect or random noise. In practice, extreme outlier expression values, as occasionally occurs in single-cell RNA-sequencing data, can distort the estimates 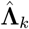. One alternative approach is to fit robust PCA^69^ instead, which incorporates a sparse matrix of large values that can model this outlier expression. While this provides a natural description of the data and can work well in some cases, we find that existing implementations of robust PCA are too slow to be applied at our scale.

Instead, we use the following fast, heuristic procedure to address the presence of outliers. After estimating 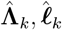 from ***R***_*k*_, we compute the median absolute deviation (MAD) of the entries in 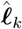 as

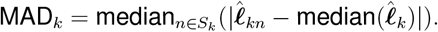

Then we define an outlier as any cell *n* where

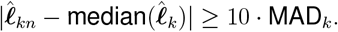

If there are any outliers, we remove the corresponding columns from ***R***_*k*_ and re-estimate 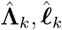. We repeat this process of outlier identification and removal until there are no outliers left, or until there are fewer than 10 cells remaining. For each estimate of **Λ**_*k*_ produced in this process, we use the absolute correlation between 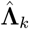 and the Wilcoxon rank-sum test statistics described in the previous section as a quality measure. We then select the 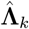 which yields the maximum absolute correlation as our final estimate.

#### Quantifying uncertainty in fingerprint estimates

To quantify uncertainty in our fingerprint estimates 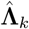, we estimate the standard error on each entry 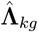. Under our model, the residual expression of gene *g* across all cells is modeled as,

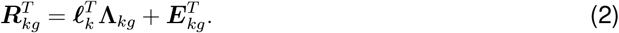

In principle, one could use Bayesian inference to quantify uncertainty under this model, but this would require high computational cost. Instead, we note that if we treat our PCA-based estimates 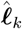 as fixed, then under Equation (2), this corresponds to ordinary least squares (OLS) regression with ***R***_*kg*_ as the response and 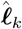 as the covariates. This motivates using the OLS estimator

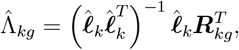

and its standard error 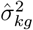 given by

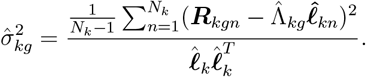

If outliers were removed to produce the estimates of 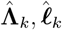, we exclude the corresponding cells from the uncertainty quantification.

#### Filtering low-signal fingerprints

If a given perturbation has minimal-to-no impact on the transcriptome, then we expect the corresponding fingerprint to contain minimal information. We filter out such low-signal fingerprints if there is likely to be little to no classification ability. This step is optional and is provided as an argument to the `LearnDictionary` function in the R package. However, we applied it with the same default parameters throughout this work and found it to improve results by limiting the contribution of noise from potentially large numbers of irrelevant fingerprints.

We assess informativeness with two metrics. The first metric is the number of the top 10 entries in 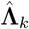, by absolute value, whose corresponding genes are marginally significant with *p <* 0.05 when comparing ***R***_0_ and ***R***_*k*_ as described under “Parameter estimation”. If a given perturbation induces transcriptional changes, then we expect the top entries of 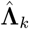 to overlap with differentially expressed genes; otherwise, the top entries are likely to just reflect noise. For a fingerprint to be retained, we require that at least two of the top 10 entries correspond to differentially expressed genes.

The second metric is computed by first projecting ***R***_0_ onto the fingerprint estimate 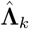. For high-signal perturbations, we expect the resulting scores 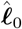 to be well-separated from the scores of the perturbed cells 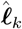. If they are not, this suggests a lack of classification ability for that perturbation. Thus, we additionally require that the marginal Wilcoxon rank-sum *p <* 0.05 between 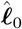 and 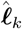.

#### Mapping query cells to fingerprints

After estimating 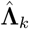 for each perturbation *k* in the reference dictionary, we next map the query data to these fingerprints (implemented in the function `FingerprintCells` in the R package). Suppose our query data consists of gene expression data with *N*^*Q*^ cells. As with the reference, we first normalize this data using SCTransform, then subset to the *G*\* genes used to estimate the fingerprints.

Let ***Y*** ^*Q*^ represent the resulting 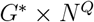 matrix of SCTransform residuals, 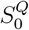 the set of 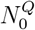 control cells for the query, 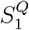 the group of 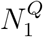 query cells to be assigned. Then

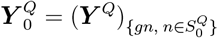

are the columns corresponding to the control cells, and

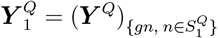

are the columns corresponding to the cells to be mapped. Here, the control cells are user-specified, and could be non-targeting controls, untreated cells, or a baseline condition. For expositional simplicity, we hereafter refer to the cells to be mapped as the “treated” cells, but these similarly depend on the biological context of the data.

We then assume the model,

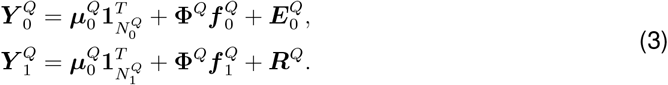

Analogously to fingerprint estimation, **Φ**^*Q*^ is a *G*\* *× M* matrix of latent factors representing shared variation, with 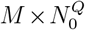 factor scores 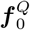 for the control cells and 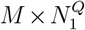 factor scores 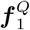 for the treated cells. These again represent the contributions to gene expression from sources of baseline heterogeneity. As before, 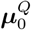 is a *G*\* *×* 1 vector of baseline mean expression in each gene, and 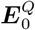 is a 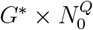 matrix of random noise.

We thus account for baseline heterogeneity by estimating 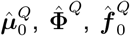, and 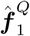 exactly as was done in fingerprint estimation. After plugging in these estimates, the remaining expression in 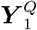, denoted as ***R***^*Q*^, can be thought of as primarily attributed to the treatment response. We model this expression as a linear combination of the fingerprints **Λ**_*k*_. Specifically, if ***r***_*n*_ denotes the *n*th cell in ***R***^*Q*^, then we assume,

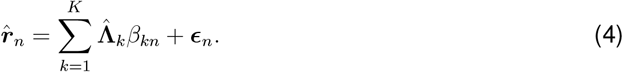

Here, ***ϵ***_*n*_ is a *G*\* *×*1 vector of random noise, and each *β*_*kn*_ represents the contribution of fingerprint *k* (to be estimated). Intuitively, fitting this regression maps cell *n* to the set of perturbations for which *β*_*kn*_s are non-zero. To fit this regression and estimate the coefficients *β*_*kn*_, we adapt the Bayesian fine-mapping method SuSiE^30^ (sum of single-effects regression) as described in the following section.

#### Sum of single-effects (SuSiE) regression

We can rewrite our regression for cell *n* in matrix notation as

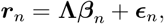

with **Λ** representing the *G*\* *×K* matrix of fingerprints and ***β***_*n*_ the *K ×*1 vector of coefficients. Then under the SuSiE model^30^,

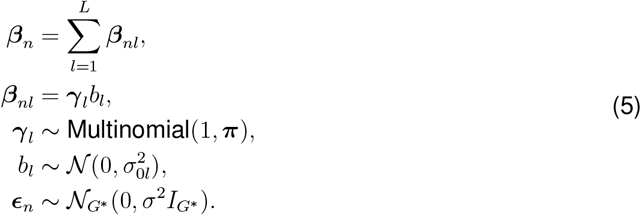

Here, the coefficient vector ***β***_*n*_ is expressed as the sum of *L* so-called single effect coefficient vectors ***β***_*nl*_, where each is constrained to only have a single non-zero entry *b*_*l*_. Intuitively, each of these single effects describes the contribution from one perturbation, and uncertainty among highly correlated fingerprints is summarized as the uncertainty in which fingerprint has the non-zero entry. This leads to a natural framework for reporting *credible sets*, each of which corresponds to one of these single effects and describes the associated perturbation(s) *k* that have high posterior probability that ***γ***_*lk*_ = 1. By default, SuSiE is fit with up to *L* = 10 such single effects allowed, and estimation proceeds through the computationally efficient iterative Bayes stepwise selection (IBSS) algorithm^30^.

SuSiE was initially developed for genome-wide association studies, and single-cell RNA-sequencing data poses some distinct statistical challenges from these data types. We highlight two main challenges in particular. The first is that ***r***_*n*_ can sometimes have even extreme outlier expression values, which can skew the estimates of the coefficients. The second is that while SuSiE was intended to use genotype data as covariates, which have little associated error, we plug in our estimates 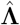, which do have associated uncertainty. To address these challenges, we developed three important extensions to the SuSiE framework, detailed in the following sections.

#### Outlier adjustment terms

First, to address the possible presence of substantially outlying expression values in ***r***_*n*_, we extend the SuSiE model to incorporate outlier adjustment terms, specifically

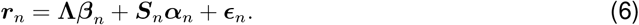

Suppose there are *A* genes *{g*_1_, …, *g*_*A*_*}* with outlying values in ***r***_*n*_. Then ***S***_*n*_ is a *G*\* *× A* matrix, where the *a*th column consists of all 0s except for a 1 in the *g*_*a*_th entry, and ***α***_*n*_ is a corresponding *A ×* 1 vector of coefficients. The intuition is that these coefficients can absorb outlier expression which might otherwise have skewed the estimates of ***β***_*n*_. As a heuristic, we set these outlying genes *{g*_1_, …, *g*_*A*_*}* as those corresponding to the top five entries in ***r***_*n*_ by absolute value, as well as any additional genes whose absolute residual values are greater than or equal to 10. The latter condition is motivated by the interpretation of SCTransform Pearson residuals as describing deviations from the expected value of each gene under the null model. We further fix the priors

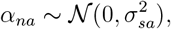

where the variances 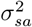 are chosen to encourage shrinkage of the *α*_*na*_ values. Specifically, we set each 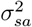 such that *P* (|*α*_*na*_| *≤* 0.5 *· r*_*na*_) = 0.95 under the prior.

To estimate the parameters of our extended SuSiE model, we leverage the IBSS algorithm. Although SuSiE was implemented for the specific model in eq. 5, the original work also presented a more general version of IBSS^30^ for fitting any additive effects model,

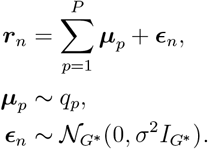

where each effect is governed by some prior *q*_*p*_.

Briefly, the IBSS algorithm iteratively updates the posterior distribution of each effect ***µ***_*p*_ while holding all other effects 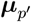 for *p*^*′*^ ≠ *p* fixed. In our setting, each effect ***µ***_*p*_ for *p* = 1, …, *L* corresponds to the single effect **Λ*β***_*np*_ as in eq. 5, and its posterior distribution and prior variance 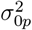 are updated exactly as in the original SuSiE implementation. Then each remaining effect ***µ***_*p*_ for *p* = *L* + 1, …, *L* + *A* corresponds to the outlier adjustment term ***S***_*n,p−L*_*α*_*n,p−L*_. Its posterior distribution can be updated analogously as a single effect with one covariate. The residual variance *σ*^2^ is updated in each iteration as an empirical Bayes estimate, and the prior variances for ***α***_*n*_ are fixed as discussed above. Convergence is determined when the evidence lower bound changes within a specified tolerance. We set *L* = 10 to allow the estimation of up to 10 different effects, and set the prior probability of inclusion 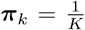 for each fingerprint in each effect. All other implementation details are as presented in Wang et al..

#### Computing credible sets

Credible sets describe the set of perturbations with high posterior probability for a given single effect. Specifically, a 95% credible set can be interpreted as a set of perturbations where the probability of containing at least one true effect is greater than or equal to 95%. Under the SuSiE framework, these are identified for each single effect *l* based on the posterior inclusion probabilities (PIPs) for each fingerprint. Each PIP

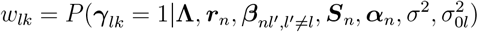

represents the posterior probability that perturbation *k* is the one associated with the non-zero coefficient in effect *l*, and is computed as in SuSiE^30^.

However, these probabilities are found conditional on the fingerprint estimates 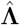, which have associated noise that is not being accounted for. To propagate this additional uncertainty, we can estimate PIPs with 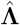 marginalized out, which we denote as 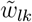. This is approximated by sampling the fingerprints according to their estimated distributions

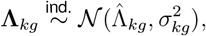

re-computing the PIPs, and averaging these results. By default, we use a total of 50 samples.

Following SuSiE, we can subsequently identify 95% credible sets for each effect *l* by ranking fingerprints in decreasing order of their PIPs 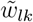 and taking the smallest set such that the cumulative sum of their PIPs exceeds 0.95. This can be interpreted as the smallest possible 95% credible set. We rank the resulting credible sets for each single effect by their associated Bayes factors, which compare the posterior probability of the perturbation effect being present to that of a null model of no effect.

In SuSiE, credible sets are filtered out either if the estimated coefficient variance 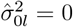 or if the credible set is not sufficiently “pure,” as assessed by the minimum absolute correlation of any two members of the credible set. This purity criterion is intended to eliminate credible sets with overly diffuse PIPs and therefore limited inferential value. We also use these two criteria, but adjust the purity criterion to instead require that the median absolute correlation be greater than 0.40. We additionally drop any credible sets whose associated log Bayes factor is less than 10% of the maximum log Bayes factor. Thus, the final output can consist of any number of credible sets from 0 (which we interpret as indicating a lack of assignment to any perturbations) to *L* = 10.

#### Proposing joint fingerprints

The resampling procedure described above propagates uncertainty from the 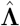 estimates into the PIPs, which results in more calibrated credible sets. However, because this procedure is applied to each single effect conditional on all the other single effects, it does not account for the impact of noise on the initial allocation of signal across effects. In some cases, a set of fingerprints might represent very similar signal, but their estimates might only be weakly correlated due to noise-driven variation and end up attributed to different effects from one another. This can be exacerbated when the signal is particularly subtle, and/or if there are low cell counts for some or all of these perturbations.

Such cases represent an opportunity to borrow strength across related perturbations. We therefore introduce a strategy to propose so-called joint fingerprints that model the shared signal across multiple perturbations, then evaluate if this joint fingerprint better explains the data than the multiple separate effects. To motivate this idea, suppose we find *J* related fingerprints *{k*_1_, …, *k*_*J*_*}* allocated across two or more separate credible sets when we might have expected all *J* to have posterior mass on a single effect. Later, we will describe how these proposals are made in an automated fashion. We assume that each of these fingerprints 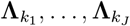 was drawn independently from a common distribution with mean 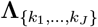. Then we can estimate

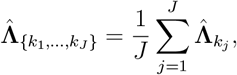

and accordingly the variance of the *g*th entry of this estimate can be set as

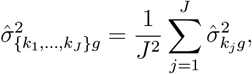

where 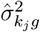 is plugged in from our earlier estimates of variance for each fingerprint.

We interpret 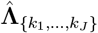 as representing the shared signal across these *J* perturbations. We then re-fit SuSiE with 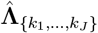alongside all the other 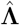 estimates, including the constituent perturbations 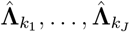, and we identify credible sets as previously described.

The model fit thus evaluates which is the better description of the data. For example, if just one credible set is found containing the joint fingerprint, we can interpret that the shared signal is the best explanation of the response, and that at least one of the *J* underlying perturbations is likely to be the true effect. In such cases, we accordingly report the final credible set as containing all of the *J* perturbations. However, if the *J* perturbations truly contributed multiple distinct effects to the response, then we would expect the PIP for 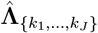 to be near-zero in every single effect, and the final result would likely be identical to the model fit with no joint fingerprint.

To identify whether a joint fingerprint should be proposed in an automated manner, we use the following procedure. We initialize perturbation sets as the credible sets, ranked by Bayes factor, from the first SuSiE fit. If there are 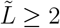 such sets, we compute the median absolute correlation between each perturbation fingerprint in set 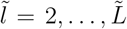 and each perturbation in the first set. If this value exceeds a threshold of 0.1, we merge the sets; otherwise, we proceed through any remaining sets up to set 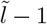. At the end of this process, if any perturbation sets were the result of combining two or more original credible sets, we propose those sets as joint fingerprints. We emphasize that any proposed joint fingerprints are statistically evaluated as described above, and thus any that are not well supported by the data are unlikely to be incorporated.

The final credible sets are identified from the SuSiE model fit with the joint fingerprints, through the same procedure described in the previous section. Within each credible set, perturbations are ranked by their PIPs. If the perturbations were from a joint fingerprint, they are ranked by the Bayes factors of the credible sets they originally appeared in, then by PIP within those sets. Occasionally, there may be multiple credible sets that overlap in the perturbations they contain (for example, a high-Bayes-factor credible set that represented the joint fingerprint, and a low-Bayes-factor credible set containing one of the constituent fingerprints). For improved interpretability, we discard any credible sets that repeat perturbations from higher-ranked credible sets. Analogously, if a perturbation appears twice within one credible set (for example, if a credible set contains both a joint fingerprint and a constituent fingerprint), we only report the perturbation once in its highest ranking position.

#### Group-level assignments

We also offer the opportunity to make assignments at the *group-level*, rather than at the cell-level. This can be desirable for two reasons. First, this can greatly improve mapping of subtle signals in particular by pooling information across cells. Second, it can be easier to interpret a single assignment for a given group (e.g. sample, condition, cluster, etc.) rather than summarizing many individual cell-level assignments.

Suppose we want to make a group-level assignment for a set of query cells in 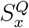. Following eq. 4,

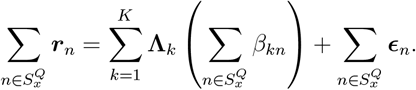

This then motivates a simple and computationally efficient strategy, which is to sum the residual expression ***r***_*n*_ for the cells in 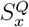 and then input the result into our framework. Under our model, there should be non-zero coefficients for every fingerprint *k* that has a non-zero coefficient in at least one cell in 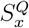.

We apply our framework exactly as described in the preceding sections, with one exception. Because extreme outlying gene expression is much less likely in aggregated profiles such as these, we use the original formulation of the SuSiE model without the outlier adjustment terms. All other steps are as indicated above. We emphasize that although this approach enables group-level classifications, every cell is still first individually processed through our latent factor model, and thus classification still benefits from modeling at single-cell resolution.

#### Computational speed-ups

Our framework naturally enables several computational speed-ups, which improves its runtime particularly on large-scale datasets. First, after estimating the baseline factors **Φ**, each fingerprint is estimated completely independently, which allows for parallelization across perturbations. Similarly, each query cell or group is also assigned completely independently, which can be parallelized as well. Second, if the initial model fit for a query cell or group does not produce any credible sets, prior to resampling the fingerprints, that would pass filtering, we just return the “unassigned” label and do not proceed with resampling.

Finally, we optionally allow the number of fingerprints inputted into the regression for each query cell or group to be capped, e.g. the top 100 fingerprints by correlation with ***r*** _***n***_ or 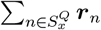 following baseline subtraction. This is conceptually related to sure independence screening^70^, which also uses marginal correlations with the response variable to reduce the number of predictors in the ultra-high-dimensional setting. In practice, we have found minimal performance differences with and without this cap, but there are substantial computational benefits. This computational speed-up is enabled in the software package, but for the analyses in this paper, we apply it only for the Genome-Wide Perturb Seq benchmarks (Figure 4).

### Statistical testing, parameters, and preprocessing

For all analyses in this paper, gene expression data was normalized using SCTransform v2^65^. For all differential expression analyses, we use the Wilcox test as implemented in the `FindMarkers` function in Seurat v5.0.3 with default parameters. We list all parameter settings for RNA fingerprinting in our description of the methods above. These values were set as default parameters in the software package, and we used these default values for all analyses in this study.

### Benchmarking on perturbation screens

#### CPA-Perturb-seq analysis

The CPA-Perturb-seq data^34^ can be accessed on GEO via accession number GSE269600. Following the original study, the hashtag oligo (HTO) counts and guide RNA (sgRNA) counts were normalized using the centered log-ratio transformation, then demultiplexed using the implementation of the MULTIseqDemux function^71^ in Seurat. Cells were retained if they had unique HTOs and sgRNAs. We then further filtered cells to those with less than 15% mitochondrial content and more than 100 total sgRNA counts. Mixscape^28^ was run using default parameters. Finally, we separately normalized the gene expression of each replicate within each cell line using the SCTransform function in Seurat.

We estimated fingerprints for each perturbation from the HEK293FT Replicate 1 cells, then mapped the HEK293FT Replicate 2 data at both the group-level (by perturbation) and at the celllevel. The set of *G*\* genes used for fingerprint estimation was chosen by performing differential expression comparing the cells of each perturbation to the non-targeting cells in Replicate 1. We took the union of the top 500 differentially expressed genes (or all marginally significant genes, if there were fewer than 500) across perturbations. We repeated the same process for the cross cell line analysis of HEK and K562 cells.

We show results for all 42 perturbations in Supplementary Figure 1C and Supplementary Figure 2B. In Figure 2B and Figure 2G, we focus on the 33 perturbations with transcriptional effects, which we define as perturbations which were neither filtered out in the original study nor all labeled as non-perturbed under Mixscape analysis.

#### CaRPool-seq analysis

The CaRPool-seq data^35^ can be accessed on GEO via accession number GSE213957. We normalized the data with SCTransform, and did not apply any additional filtering steps. For each single-gene perturbation, we randomly withheld 50% of the cells. Using only the non-withheld cells, we applied the same feature selection strategy as in the CPA-Perturb-seq analysis and learned fingerprints for each single-gene perturbation. We then mapped the withheld cells with single-gene perturbations at the group-level (by perturbation), considering only those that were not filtered out during fingerprint estimation. The 11 perturbations assigned with log Bayes factor greater than 800 were considered to have sufficient classification power for the dual perturbation analysis. We then mapped all corresponding dual perturbations to the subset of the dictionary containing these 11 perturbations at both the group and the single-cell level. For Figure 3C, any of those dual perturbations whose constituent perturbation fingerprints had a correlation of less than 0.1 was considered to be comprised of uncorrelated effects.

#### Genome-Wide Perturb-Seq analysis 1

The processed Genome-Wide Perturb-Seq (GWPS) datasets^19^ were obtained from Figshare with accession number 20029387 and normalized with SCTransform. We estimated fingerprints from the main genome-scale GWPS dataset, then mapped data from the K562 essential-scale screen at both the group-level (by perturbation) and individual cell-level. Because of the large number of perturbations, instead of performing differential expression, we used all features in the processed GWPS data, which consists of all genes expressed at greater than 0.01 UMI per cell. When mapping, we subsetted the K562 essential-scale screen to just perturbations that were not filtered out in fingerprint estimation. We further randomly held out 100 non-targeting cells to be classified alongside the perturbed cells. As described in the “Computational speedups” section, inspired by sure independence screening^70^, we internally capped the number of fingerprints inputted into each SuSiE regression to the top 100 most correlated fingerprints for each group or cell, following processing via our latent variable model, due to the large data size and number of classifications.

We also repeated these steps to map data in the RPE1 essential-scale dataset at the group-level. Enrichment analysis of the classifications was run using the enrichGO function from the R package clusterProfiler^72^. For the correct classifications, the background set was all perturbations which were profiled in the RPE1 essential scale experiment, and also correctly classified (i.e. defined as having the correct perturbation in the top credible set) in the K562 essential-scale data. For the incorrect classifications, the background set was the same, and the input set was the subset that was either unassigned in the RPE1 data, or incorrectly assigned such that the maximum absolute correlation between the true fingerprint and any fingerprint in the top credible set was less than 0.1.

#### Long Island City plots

We introduce Long Island City plots, inspired by Manhattan plots in genome-wide association studies, to summarize RNA fingerprinting perturbation assignments. In each plot, the x-axis consists of the set of fingerprints in the dictionary, ordered by hierarchical clustering of the fingerprint correlation matrix. In Manhattan plots, the x-axis denotes the fixed order genomic location, which naturally results in correlated SNPs being close together in the plot. While our ordering has no direct positional analogy, we cluster so that correlated signals also localize on the x-axis. The y-axis represents the posterior coefficient for each fingerprint, conditional on inclusion, under our model fit using all available fingerprints. For each fingerprint, the coefficient is chosen as the one for the single effect where that fingerprint’s Bayes factor is largest. Note that in the full method, there can be a second model fit using joint fingerprint proposals (see “Proposing joint fingerprints”), which this visualization does not depict. In our R package, this visualization is implemented in the function `LongIslandCityPlot`.

#### Bayes factor analysis for gene identification

To identify the genes that contributed the most to a particular assignment, we used our initial model fit based on all available fingerprints and without joint fingerprint proposals. We identified the credible set where the Bayes factor for the assignment of interest was largest, then systematically re-computed this Bayes factor with each gene removed. The genes resulting in the largest changes to the Bayes factor are considered to be the most important. In our package, this step is implemented in the function `ExplainMatch`.

### Comparisons to other methods

To compare our performance against other methods, we used a subset of the GWPS benchmark data, because elastic net regression was not able to scale to the full dataset. Specifically, we used a randomly selected set of 1,234 perturbations from GWPS (one-eighth of the total dataset) and estimated fingerprints, of which 525 remained after filtering. We then mapped individual cells from the subset of the K562 essential-scale data that belonged to these 525 perturbations. The same features were used as when mapping the K562 essential-scale data in the full benchmark described earlier. For each comparator method, detailed in the subsections below, we only supplied the subset of the GWPS data with these 525 perturbations for training, and the same query data that we mapped was given for testing.

#### Elastic net regression

We used the implementation in the R package glmnet to fit an elastic net multinomial regression model to predict perturbation labels from the gene expression profiles. Specifically, we fit the model on the SCTransform residuals of the training data described above, subset to the same features we used for fingerprint estimation, with the target gene identity (including non-targeting) as the response variable. We used 10-fold cross-validation and set the *α* parameter to 0.5. To make predictions on the query data, we used the value of the regularization parameter *λ* that yielded the minimum mean cross-validation error. We assigned each cell the label that corresponded to the highest predicted probability, which could include the non-targeting label.

#### GEARS

We ran GEARS^36^ following the online tutorial to predict a response vector for each perturbation. Specifically, we applied the recommended preprocessing steps to the raw counts of the training data and chose the top 5, 000 highly variable features as suggested. We trained the model using hyperparameters set to the same values as in the tutorial, except we increased the number of epochs to 20. After training, we then predicted the perturbation responses.

We assessed whether these predicted transcriptomic profiles could be repurposed to enable matching, since matching to query data is not an inherent functionality of GEARS. Specifically, we first processed the raw counts of the query data using the same preprocessing steps. For each non-targeting cell, we computed its maximum correlation with any predicted perturbation response, and then used this resulting distribution of correlations as an empirical null distribution.

For each perturbed cell, we identified the perturbation response with the highest correlation, and assigned its label to that cell if the correlation exceeded the 95th quantile of the empirical null distribution. Otherwise, the cell was labeled as “unassigned.”

#### CPA

We ran CPA^37^ following the online tutorial for context transfer to predict a response vector for each perturbation. We applied the recommended filtering and preprocessing steps to the counts of the training data, including selecting the top 5, 000 highly variable genes. We then trained the model using the same hyperparameters as in the tutorial. We preprocessed the counts of the non-targeting cells in the query data using the same steps, then applied the CPA model to predict what each of the 525 perturbation responses would look like in those cells. We used the mean of the resulting log-transformed responses to summarize each predicted perturbation response. These response summaries were then used to label each perturbed query cell as was done for GEARS above.

#### F1 scores

F1 scores were computed for each perturbation under each approach as the harmonic mean of the sensitivity and the precision. The sensitivity was computed as the fraction of cells of a given perturbation whose top label was correct, and the precision was computed as the fraction of cells whose top label was a given perturbation that actually received that perturbation. A cell that was labeled “unassigned” was considered incorrect. Scores were left undefined for any given perturbation and approach if the precision could not be computed, i.e. if no cells at all were assigned a particular perturbation label. For RNA fingerprinting, the label used for each cell was the top member of the top credible set.

#### Altered target gene expression

For RNA fingerprinting and elastic net regression, we additionally mapped cells from an altered version of the query dataset, with no changes to the fingerprints in our approach or to the trained model in elastic net regression. Specifically, for each perturbation, we set the SCTransform residuals of the target gene for cells receiving that perturbation to 0.

### Ribosomal protein analysis

#### Fingerprint estimation

To explore structure among the RPL perturbations in the RPE1 essential-scale dataset, we estimated fingerprints for each perturbation using the same set of features as when mapping to GWPS. We then ran PCA on the resulting matrix of fingerprints using the irlba package.

#### Identification of gene modules

To identify a shared module of genes among ribosomal perturbations in the RPE1 data, we performed differential expression comparing all cells that received an RPL or RPS perturbation to the non-targeting cells, and retained genes with an adjusted p-value of less than 0.05. To quantify expression of this shared module for each perturbation, we first ran Mixscape^28^ on the cells with RPL perturbations and filtered out any cells labeled as non-perturbed. We then used the AddModuleScore function in Seurat to compute two scores in each remaining cell: one using all the down-regulated genes from the shared module, and one using all the up-regulated genes. We defined the overall shared module expression in each cell as the difference between these two scores, and summarized each perturbation with the median of this value over all its cells. In subsequent analyses focusing only on RPL perturbations that successfully induced ribosomal stress (Figure 5F, Supplementary Figures 5A and B), we excluded perturbations where this score was less than 0.1. To identify a distinct module for our four RPL perturbations of interest, we defined two groups of RPLs based on our PCA analysis (Figure 5B): RPL5/11/10/24 and RPL39/26/7L1/37. We performed differential expression of cells within these two groups, and retained all genes with an adjusted p-value of less than 0.05.

#### Bulk RNA-sequencing data of nutlin-3 treatment

The bulk RNA-sequencing data for nutlin-3-treated RPE1 cells^45^ was obtained from GEO under accession number GSE86104. We processed this data through the standard log-normalization and scaling pipeline in Seurat.

#### dsiRNA experiment

##### Cell culture and siRNA knockdown

A549 cells were maintained in DMEM media (Caisson Labs: DML10) supplemented with 10% FBS (Corning:35-010-CV) and 1X Non-essential amino acids (Sigma: M7145-100ML). All cells were grown at 37°C and 5% CO_2_. 30,000 cells were plated in each well of a 24-well plate in 500 *µ*l of media. 34 hours later, 12 pmols of siRNA were transfected with lipofectamine for a final concentration of 20 nM in each well. The sequences of each siRNA are available in Supplementary Table 1. The next day, 24 hours later, each well was treated with either DMSO (vehicle control) or 5 *µ*M nutlin-3a (Selleck Chemicals: S8059). The cells were left in treatment for 24 hours before harvesting.

##### FlexPlex profiling

Cells were profiled using the FlexPlex protocol described under the “FlexPlex experimental protocol” and “FlexPlex computational processing” subsections. From the resulting data, we first normalized the Hashtag-derived Oligo (HTO) counts with the centered log-ratio transform, demultiplexed with `MULTIseqDemux`, then filtered out any cells without a unique HTO assignment. We then normalized the gene expression data with SCTransform. A fingerprint was estimated for nutlin-3a treatment from the non-targeting cells, where the DMSO non-targeting cells were used as the control condition. Features with a mean UMI count of at least 0.1 across all cells were included.

##### Western blot

Cells were harvested in lysis buffer containing 2 mM EDTA (Sigma: E5134), 2 mM EGTA (Sigma: E3889), 1% Triton-X (Sigma: T8787), and 0.5% SDS (Sigma: 71736) in 1x PBS with EZ Block phosphatase inhibitor cocktail (BioVision: K273-1) and Complete mini EDTA-free protease inhibitor cocktail (Sigma: 4693159001). Total protein was determined by BCA assay (ThermoFisher: 23227) and 15 *µ*g of protein in Nupage LDS sample buffer (Thermo: NP0007) were loaded onto Novex 4-12% Tris-Glycine minigels (Thermo: XP04125). Proteins were transferred to PVDF membranes (EMD Millipore: IPFL07810), blocked in 5% milk in TBS-Tween for one hour at room temperature (RT), and incubated with primary antibodies diluted in 5% milk in TBS-Tween overnight at 4°C. The following day, membranes were incubated with HRP-conjugated secondary antibodies (Jackson ImmunoResearch: 111-035-003;115-035-003) for one hour at RT, washed, incubated with chemiluminesence substrate (Thermo: 34577) and scanned on a Licor C-digit device. The concentration and catalog number of primary antibodies were as follows: p53 1:1000 (Cell Signaling: 9282T) and Histone H3 1:2000 (Cell Signaling: 4499). 1

### Mapping drug mechanism of action

#### Sci-Plex positive control analysis

The sci-Plex data^20^ can be accessed from GEO at accession number GSM4150378. We used the data in K562 cells following 24 hours of treatment, and normalized these data with SCTransform without any additional filtering. We estimated fingerprints from GWPS using the intersection of genes with expression greater than 0.01 UMI per cell in GWPS, and genes present in the sci-Plex dataset, then mapped sci-Plex drug-dose combinations at the group-level. The untreated, vehicle-only cells were used as the control condition during mapping.

#### Alternative drug-target analyses

We applied three comparative approaches to assess the identification of the expected target in GWPS from three of the sci-Plex drug-dose combinations (Dasatinib 100 nM, Flavopiridol 10 nM, and GSK-LSD1 2HCl 10 nM).

First, we conducted a gene set enrichment overlap test (Supplementary Figure 6B-D) by using Fisher’s test to evaluate significant overlap between the differentially expressed genes from the drug-dose combination, compared to vehicle control, and the differentially expressed genes from each genetic knockdown in GWPS, compared to non-targeting. Differentially expressed genes were determined as those with an adjusted p-value of less than 0.05. The background gene set consisted of all features used in computing fingerprints, and we only considered GWPS knockdowns corresponding to fingerprints that passed filtering.

Second, we conducted a rank-based gene set enrichment test (Supplementary Figure 6B-D) by comparing the differentially expressed genes from each drug-dose combination to the genes ranked by absolute log fold-change, as computed by Seurat, under each genetic knockdown. The same features and perturbations were considered as above. The enrichment test was performed as implemented in the R package fgsea^73^.

Finally, we compared each drug-dose combination to each genetic knockdown, using the same features and perturbations as above, via cosine similarity (Supplementary Figure 6B-D). Within the sci-Plex and GWPS datasets respectively, we pseudobulked gene expression by condition (drug-dose combination or perturbation), then log-normalized and scaled as implemented in the default Seurat pipeline. We then computed cosine similarity on these scaled profiles.

#### HDAC perturbation screen experimental details

##### Cell culture

K562-CRISPRi-v2 cells were derived as in previous work^29^ and were maintained in IMDM media (ThermoFisher, 31980030) supplemented with 10% FBS (Corning, 35-010-CV) and 1 mM Pen-Strep (Sigma, P0781-50ML). HEK293FT cells were acquired from Thermo Fisher (R70007) and maintained in DMEM media (Cytiva, SH30243.01) supplemented with 10% FBS. All cells were grown at 37°C and 5% CO_2_.

##### CRISPR guide library cloning

One guide pool was designed using four guides per gene: two guides from the Genome-Wide Perturb-seq CRISPRi guide library^19^ and two guides from the Dolcetto CRISPRi guide library^74^ (Supplementary Table 2). A partial hU6 promoter sequence (GGAAAGGACGAAACACCG) and partial guide scaffold overhang (GTTTAAGAGCTAAGCTGGAAACAGC) were added to each guide sequence and were ordered as an oligo pool (oPool) from Integrated DNA Technologies with the following format:

Partial-hU6-promoter :: gRNA :: partial-guidescaffold.

The oligonucleotide pool was amplified for cloning with the following primers:

- Forward: 5’-TAACTTGAAAGTATTTCGATTTCTTGGCTTTATATATCTTGTGGAAAGGACGAAACACCG-3’
- Reverse: 5’-GACTAGCCTTATTTAAACTTGCTATGCTGTTTCCAGCTTAGCTCTTAAAC-3’

50 ng of pooled oligonucleotides were amplified with the above primers and KAPA HiFi Hot-Start ReadyMix (Roche, 07958935001) using the following PCR program: 98°C for 30 seconds, three cycles of 98°C for 10 seconds, 60°C for 15 seconds, and 72°C for 15 seconds, followed by 72°C for 3 minutes. The PCR product was purified using a 2.0x SPRI cleanup. The amplified pool was ligated into a digested modified CROP-seq cloning vector^75^ using a Gibson Assembly reaction (NEB, E2611S), following the manufacturer’s recommendations.

Ligated plasmids were concentrated through isopropanol precipitation and resuspended in TE buffer. Concentrated plasmid libraries were electroporated into Endura electrocompetent cells (Lucigen, 60242-1), recovered in warm SOC media for 1 hour, and plated onto ampicillin containing LB-Agar plates. Bacterial colonies were harvested into Luria Broth and purified using a Maxiprep kit (IBI Scientific, IB47121). Guide RNA balance was verified through sequencing of the plasmid library on an Illumina MiSeq instrument.

##### Lentiviral production

11 million HEK293FT cells were seeded in a 10 cm dish 18 hours before transfection. The transfection reaction used 45 *µ*l, 1 mg/ml polyethyleneimine (PEI; Polysciences 23966-1) and 20 *µ*g total plasmid DNA (6.4 *µ*g psPAX2: Addgene, 12260; 4.4 *µ*g pMD2.G: Addgene, 12259; 9.2 *µ*g sgRNA-expressing plasmid). Six to eight hours post-transfection, the medium was exchanged for 10 ml DMEM and 10% FBS containing 1% bovine serum albumin (VWR 97061-420). Viral supernatants were collected after an additional 48 hours, spun down to remove cellular debris for 5 minutes at 4°C and 1,000 g, passed through a 0.22 *µ*m filter, and stored at −80°C until use.

##### Lentivirus transduction

K562 CRISPRi cells were transduced at a low multiplicity of infection (MOI) (0.05-0.1) with lentivirus containing the guide pool. For 8 days of selection for guide positive cells, the K562 cells were cultured with 2 *µ*g ml^*−*1^ puromycin (Thermo Scientific, A1113803) and culture expansion and media changes every three days. On day 8, dead cells were removed from the culture with a dead cell removal kit and columns (Miltenyi Biotec: 130-090-101; 130-042-401) and live cells were collected.

##### Direct capture Perturb-seq and sequencing

We performed scRNA-seq (10x Genomics Chromium Single Cell 3’ Gene Expression v3 with Feature Barcoding technology for CRISPR screening). Gene expression and sgRNA feature libraries were constructed following the manufacturer’s protocol. The libraries were quantified using a qubit and run on a NextSeq 550 instrument. Data were processed using Cell Ranger.

#### FlexPlex drug screen experimental details

##### Cell culture

K562 were maintained in IMDM media (ThermoFisher: 31980030) supplemented with 10% FBS (Corning:35-010-CV), 1 mM Pen-Strep (Sigma: P0781-50ML) and 1X Non-essential amino acids (Sigma:M7145-100ML). HEK cells were maintained in DMEM media (Caisson Labs: DML10) supplemented with 10% FBS (Corning:35-010-CV) and 1X Non-essential amino acids (Sigma: M7145-100ML). All cells were grown at 37°C and 5% CO_2_.

##### FlexPlex experimental protocol

In addition to the summary below, we have published a detailed protocol for FlexPlex at https://phospho-seq.com/post/protocol6/FlexPlexProtocol.pdf. We also include probe, hashtag oligo, and antibody-derived tag sequences in Supplementary Table 3.

For each condition in each experiment, up to two million cells were harvested from growth media, placed into PCR strip tubes, and resuspended in 95 *µ*l of PBS. Cells were fixed by adding 6 *µ*l of 16% formaldehyde (Sigma: F8775-25ML) (final concentration, 1% FA in PBS). Cell suspensions were left to fix for 10 minutes at RT, with inversion every 3 minutes. The fixation reaction was quenched by adding 13.7 *µ*l 1M glycine and filling the tubes with ice-cold PBS. The suspensions were centrifuged for 5 minutes at 400 x g at 4°C, after which the supernatant was removed. The cells were resuspended in 200 *µ*l PBS, and the centrifugation was repeated.

After the second centrifugation, the cells were resuspended in 50 *µ*l of lysis buffer (10 mM Tris-HCl, 10 mM NaCl, 3.33 mM MgCl2, 0.1% NP-40 (Thermo: 28324), 1% BSA, 0.1 mM DTT (Invitrogen: y00147), 100U NxGen RNAse Inhibitor (Lucigen: 30281-1) in H_2_O and incubated on ice for 5 minutes for permeabilization.

After 5 minutes, 200 *µ*l of wash buffer 1 (10 mM Tris-HCl, 10 mM NaCl, 3.33 mM MgCl2, 1% BSA, 0.1 mM DTT, 20U NxGen RNAse Inhibitor in H_2_O) was added, and cells were centrifuged for 5 minutes at 500 x g after which the supernatant was discarded and the cells were resuspended in 100 *µ*l blocking buffer per hashing group and placed in a tube rotator at RT for 30 minutes. Blocking buffer consisted of 100 *µ*g 30 nt blocking oligo, (`NNNNNNNNNNNNNNNNNNNNNNNNNNNNN/3ddC/`), 0.1 mM DTT, 100U Nx-Gen RNAse Inhibitor and 3% BSA in PBS. For hashing, we spiked in 15 *µ*l of 200 *µ*M unique OligoHash per hash group. OligoHashes were ordered from IDT with the following sequence: `CGGAGATGTGTATAAGAGACAGXXXXXXXXXXXXXXXGCTTTAAGGCCGGTCCTAGC*A*A` where the *s represent the unique OligoHash barcode.

While in the blocking and hashing step, the primary antibody pool was incubated with single-stranded DNA binding protein (Promega: M3011) to help reduce off-target effects^76,77^. Briefly, 8 *µ*g of SSB was added per 1 *µ*g of antibody in a solution of 1x NEB buffer 4 (NEB:B7004S) for 30 mins at 37°C. After this incubation, BSA and PBS were added to make a final solution of 3% BSA in 1X PBS. Finally, 0.1 mM DTT and 100U/100 *µ*l NxGen RNAse Inhibitor were added to the staining solution.

After blocking, cells were centrifuged at 600 x g for 5 minutes and washed with a wash buffer of 3% BSA and 0.1% Tween in PBS twice. After the second wash, the individually hashed pools were combined in wash buffer and centrifuged again before resuspending in the previously prepared primary antibody staining solution with a maximum of two million cells. The cells were placed on a tube rotator at room temperature for 1 hour for primary antibody staining. After 1 hour, the cells were centrifuged and washed three times in the same wash solution from the blocking step. After the final supernatant was removed, the cells were input into the manufacturer recommended 10x Flex workflow starting with 4% formaldehyde fixation, which acts as a second fixation step.

We used 10x Flex v1 kit for single-plexed samples with feature barcoding for the HDAC inhibitor experiment and the 10x FLEX GEM-X kit for the dsiRNA experiment. We made minor modifications to the protocol to facilitate the capture of OligoHashes, TotalSeq-B antibodies and in the siRNA experiment, custom probes. Custom RNA probes were spiked into the probe hybridization step using 10x Flex recommended concentrations. The custom probe sequences can be found in Supplementary Table 3. Next, because we used OligoHashing in each of our experiments, which enables high-throughput doublet demultiplexing^78^, we can superload the microfluidic chip over the recommended loading quantity. Next, we spike in 1 *µ*l of 0.2 *µ*M an OligoHash additive primer into the pre-amplification step, in order to preamplify the OligoHashes alongside the mRNA, guides and ADTs. The sequence for the OligoHash additive primer is: CGGAGATGTGTATAAGAGAC*A*G. Once the preamplification is finished, the final products are eluted into 100 *µ*l of elution buffer. Each modality will use 20 *µ*l of this solution as the basis for modality specific PCR. Transcriptome and ADTs are amplified according to the protocol as written. OligoHashes are amplified by using a Nextera Read 2 index primer (CAAGCAGAAGACGGCATACGAGAT NNNNNNNNGTCTCGTGGGCTCGGAGATGTGTATAAG) and SI-PCR primer (10 *µ*l of 10 *µ*M each). Each modality is amplified under the same PCR conditions as written for the transcriptome, with 12-14 cycles for ADTs and OligoHashes. Sequencing for all libraries was performed using a NextSeq 550 instrument with the read structure of 28/8/50. Optimal read numbers per cell vary, but are generally 10,000 for gene expression, 5,000 for ADT, and 2,000 for OligoHashes.

##### FlexPlex computational processing

Transcriptomic data was aligned and quantified using CellRanger 8.0 to generate count matrices. OligoHashes (HTOs) and ADTs were quantified using a custom script that scanned the FASTQ files for cell barcodes, UMIs and known index sequences, building a counts matrix from this information. Each version of the quantification script can be found at https://phospho-seq.com/files/quant.ipynb. HTOs were normalized using CLR normalization, and we performed demultiplexing and doublet detection using the `MULTIseqDemux` function with default parameters. Only cells classified as singlets were kept. ADTs were also normalized using CLR normalization. Gene expression was normalized using SCTransform.

#### Fingerprint estimation

We estimated and combined fingerprints from three different dictionaries: sci-Plex, GWPS, and the HDAC perturbation screen. The features used were the intersection of genes with expression greater than 0.01 UMI per cell in GWPS, and genes present in each other relevant dataset. We estimated fingerprints from sci-Plex and GWPS as before, with the exception that we only retained GWPS fingerprints corresponding to HDAC perturbations.

For the HDAC perturbation screen, we first normalized the guide counts with the centered log-ratio transformation, demultiplexed with `MULTIseqDemux`, then filtered out any cells that did not have a unique guide. We then normalized the gene expression data with SCTransform, and estimated fingerprints for each individual guide.

#### Trajectory analysis

We fit a “pseudo-dose” trajectory to each dose of the HDAC inhibitors in the sci-Plex data, by considering all retained fingerprints whose corresponding drugs are labeled as targeting HDAC in the original data annotations. We ran PCA with two dimensions using the irlba package, then used Slingshot^79^ to fit a trajectory in the resulting lower-dimensional space.

#### Bulk RNA-sequencing data of HDAC1/2/3 knock-out

We compared the gene expression patterns along the pseudodose trajectory to bulk RNA-sequencing data of HDAC1/2/3 triple knock-out HeLa cells^55^. These data were obtained from GEO under accession number GSE234005, then processed through the standard log-normalizatio and scaling pipeline in Seurat.

#### Mapping HDAC inhibitors

We mapped cells treated with Ricolinostat and ACY-738 from the FlexPlex data to these combined fingerprints at the group-level for each drug-dose combination. The untreated, DMSO cells were used as the control condition. Pseudodose was imputed for each drug-dose combination as the mean pseudodose over each HDAC inhibitor in the top credible set.

### Immune response analysis

#### Fingerprint estimation

To estimate fingerprints from the Immune Dictionary^22^, we chose the set of features as the genes with expression greater than 0.01 UMI per cell in the entire dataset. We then estimated fingerprints separately for each cell type, using the PBS cells as controls, and excluding cell types with fewer than 30 PBS cells available. Metadata such as cell type and cytokine labels were used as provided by the original authors. The counts data are available at GEO under accession GSE202186.

#### Mapping influenza rechallenge data

The influenza rechallenge dataset can be accessed on GEO via accession number GSE194058. Metadata such as mouse sample ID and cell type were used as provided by the original authors. The data corresponding to each major cell type (i.e., excluding the three smallest clusters, which are the alveolar macrophages, ILC2 cells, and cycling B/T cells) was mapped to the corresponding fingerprints of the same cell type from the Immune Dictionary, using the cells from the resting mice as the control condition. Mapping was done at the single-cell level.

For three such cell types, namely those labeled as “DC,” “mono/Mac,” and “neutrophils,” there was no set of fingerprints directly corresponding to the identical cell type label. For the former two, this is because the Immune Dictionary dataset clustered myeloid cells at different levels of granularity from the influenza rechallenge dataset; for the latter, this is because the Immune Dictionary contained many fewer neutrophils, and this cell type was filtered from fingerprint estimation for having too few control cells available. To identify the best match from the available fingerprints, we mapped each of those cell types at the group-level (combining across all rechallenged mice) to each set of fingerprints one at a time, and selected the fingerprints which yielded the highest log Bayes factor in the top credible set. 1

#### Characterizing B cell heterogeneity 1

B cells which received assignments were divided into clusters based on their top credible set. We noted two main groups of cytokines that tended to co-occur: the “primarily Type 1” group consisted of IFNA1, IFNB, IFNE, IFNK, and IL-7, and the “primarily IFNG-related” group consisted of IFNG, IL-15, IL-18, the combination IL-2 + IFNG, and the combination IL-2 + IL-15. If a cell’s top credible set contained at least one cytokine from both groups, it was labeled as having “both responses”; if it contained at least one cytokine from the former but not the latter, it was labeled as having a “primarily Type 1 response”; if it contained at least one cytokine from the latter but not the former, it was labeled as having a “primarily IFNG-related response”; and if, rarely, it contained some other cytokine, it was labeled as “other.”

#### Comparison to Immune Response Enrichment Analysis

To compare our results, we ran the Immune Response Enrichment Analysis module on the Immune Dictionary web app^22^ on the up-regulated differentially expressed genes (adjusted *p*-value *<* 0.05) of rechallenged B cells (both SI and SI+CLL) versus resting B cells. This was done with default parameters.

## Notes

https://github.com/satijalab/rna-fingerprinting

https://zenodo.org/doi/10.5281/zenodo.17127744

