## Supplementary Figures for "Mapping transcriptional responses to cellular perturbation dictionaries with RNA fingerprinting"

### Supplementary Materials for: Mapping transcriptional responses to cellular perturbation dictionaries with RNA fingerprinting

### Supplementary Figures

#### Supplementary Figure 1

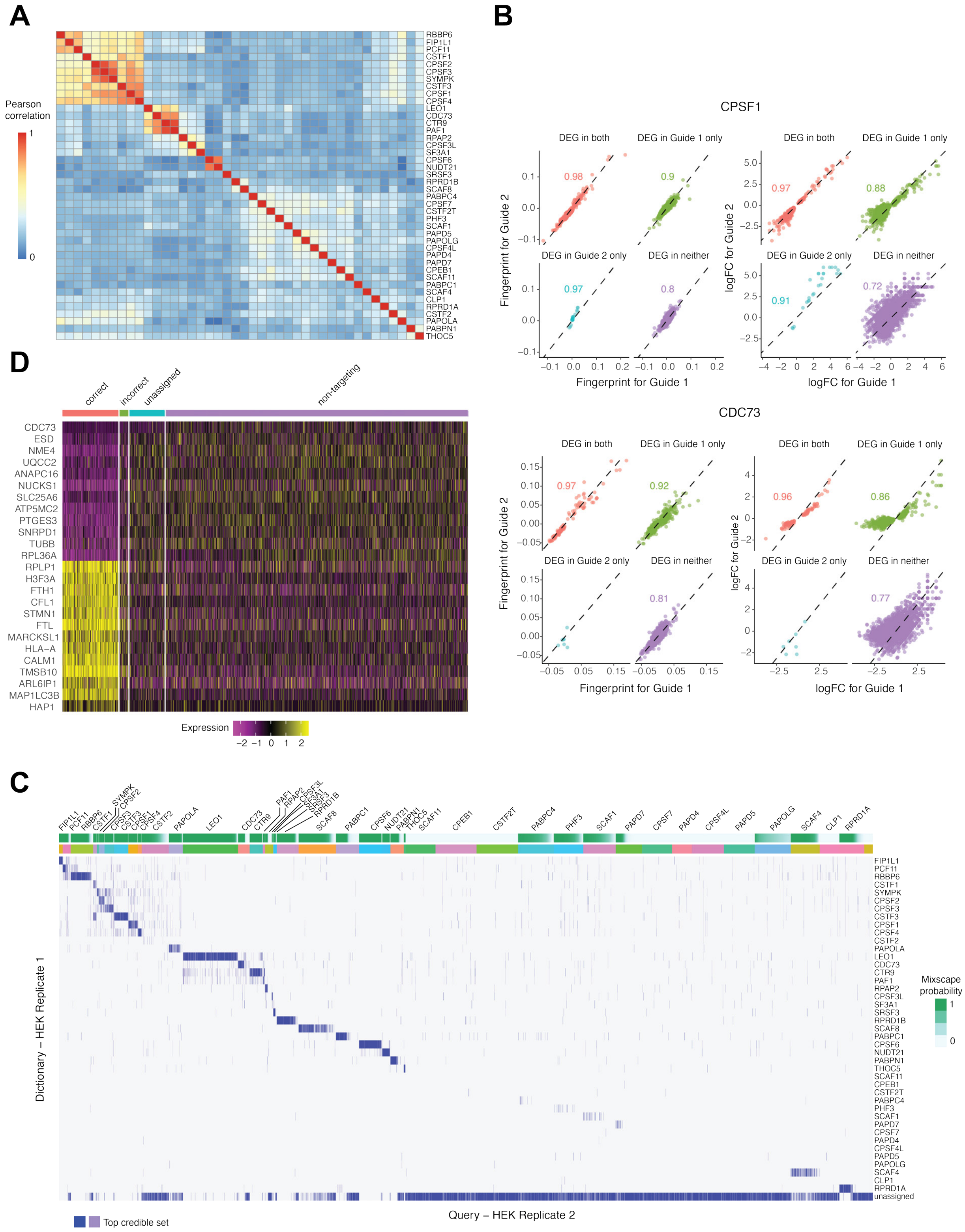

#### Supplementary Figure 1

(A) Heatmap of correlation matrix for fingerprints estimated from the CPA-Perturb-seq data in HEK cells. Fingerprints estimated for members of the same complex (e.g. CPSF6 and NUDT21) are highly correlated.
(B) Comparison of fingerprints and differential expression (DE) statistics for two independent guides targeting CPSF1 (top) and CDC73 (bottom). Left: scatterplots of fingerprint values for Guide 1 versus Guide 2, colored by genes identified as differentially expressed compared to non-targeting controls in both guides, only one guide, or neither. Right: scatterplots of $\log_2$ fold changes from DEG analysis for the same comparisons. Fingerprint estimation shows improved guide reproducibility, even for genes that exhibit subtle changes and would not be conventionally detected as differentially expressed. DE analysis was performed using Wilcox test in `Seurat` with an adjusted p-value threshold of 0.05.
(C) Heatmap of individual cell-level fingerprinting results across replicates for all perturbations. Each row corresponds to a reference fingerprint, and each column corresponds to a query cell, grouped by ground-truth labels. Shading denotes inclusion of the fingerprint in the query cell's top credible set. Blue indicates the top credible set, and dark blue indicates the top member.
(D) Heatmap of gene expression for all cells which received CDC73 guides, as well as a random sample of non-targeting cells. CDC73-targeted cells are split by whether they were assigned correctly (defined as the presence of CDC73 in the top credible set), incorrectly, or left unassigned.

### Supplementary Figure 2

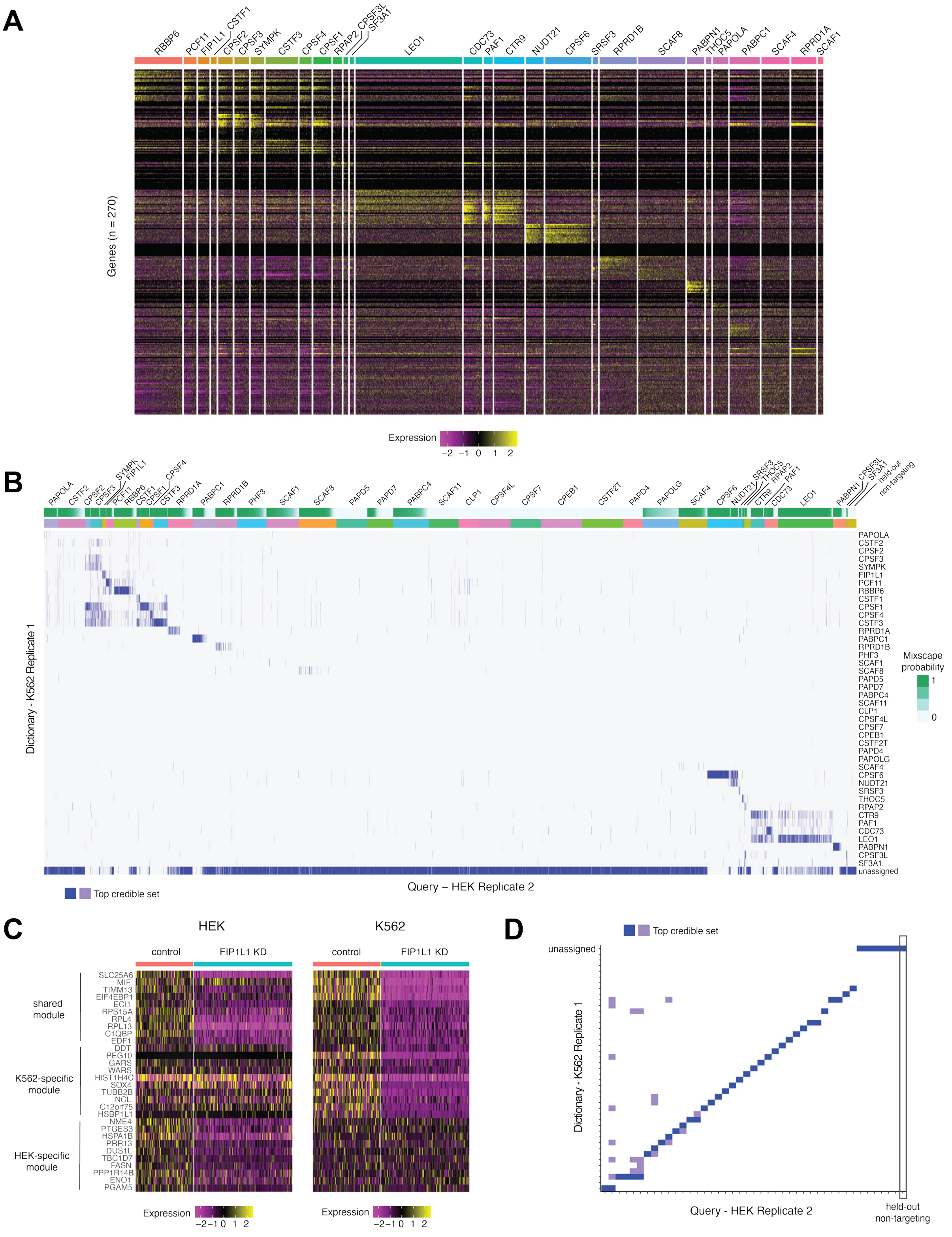

#### Supplementary Figure 2

(A) Heatmap of gene expression for query HEK cells in the CPA-Perturb-seq data, split by their assignments (summarized as the top member of the top credible set). Assignments are only shown if at least 100 cells were given that label. Unassigned cells not included.
(B) Heatmap of individual cell-level fingerprinting results across contexts for all perturbations. Each row corresponds to a reference fingerprint, and each column corresponds to a query cell, grouped by ground-truth labels. Shading denotes inclusion of the fingerprint in the query cell's top credible set. Blue indicates the top credible set, and dark blue indicates the top member.
(C) Example perturbation (FIP1L1) that exhibits divergent transcriptional responses between HEK (left) and K562 (right) cells.
(D) Group-level fingerprinting results for RNA fingerprinting across cell lines. Heatmap structure and shading are the same as in (B).

### Supplementary Figure 3

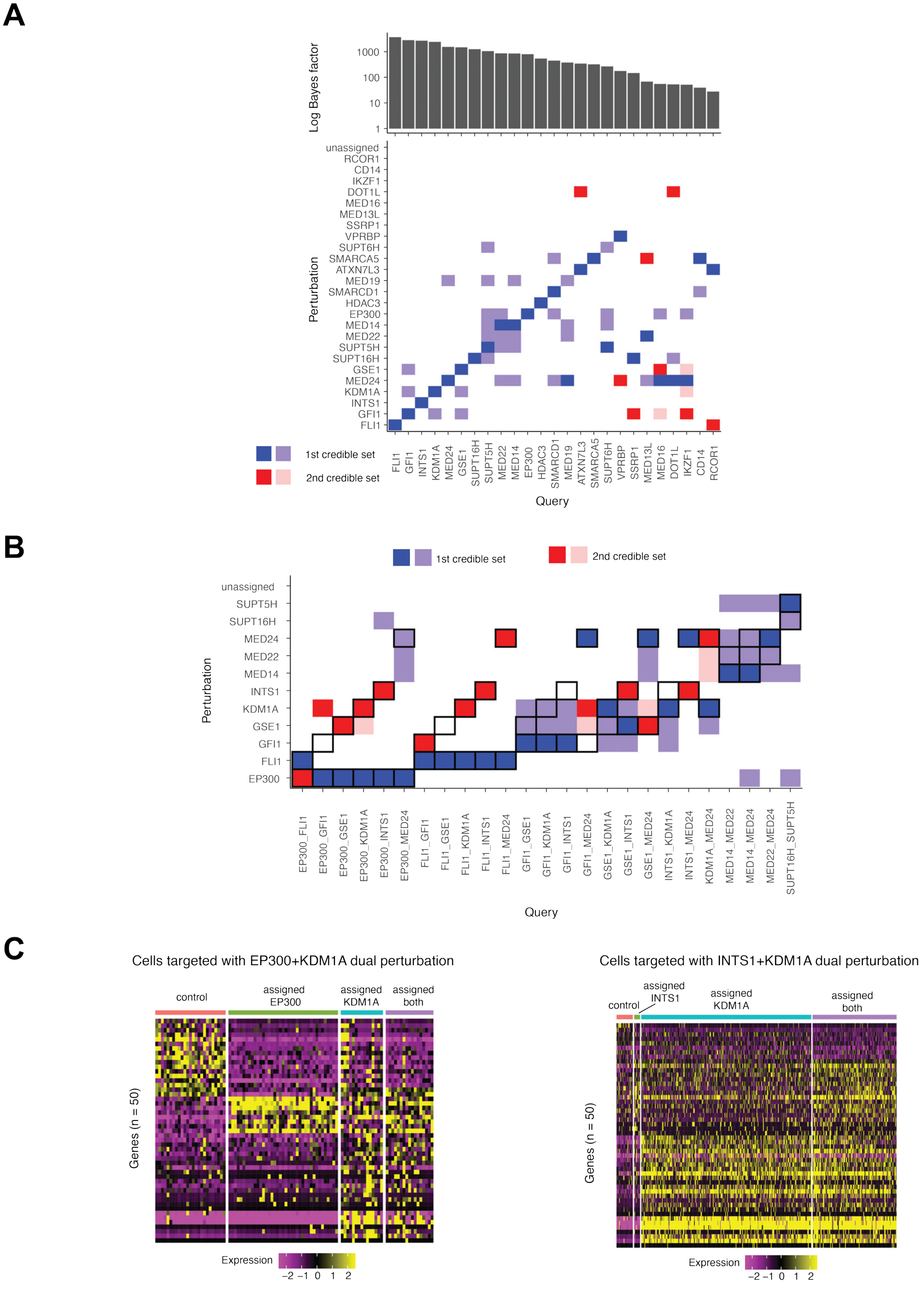

#### Supplementary Figure 3

(A) Group-level fingerprinting results for single-gene perturbations in the CaRPool-seq data. Heatmap shows the dictionary perturbations included in the first credible set (blue) or second credible set (red) for each query perturbation. Bayes factors for the top credible set are shown at the top.
(B) Group-level fingerprinting results for dual perturbations. Heatmap shows the dictionary perturbations included in the first credible set (blue) or second credible set (red) for each query perturbation. Black boxes denote the expected ground-truth results.
(C) Gene expression profiles for cells targeted by the EP300+KDM1A dual perturbation (left) or the INTS+KDM1A dual perturbation (right), grouped by assignment to each constituent perturbation, both, or unassigned. Heatmap shows that the assignment from RNA fingerprinting reflects the cell's underlying molecular phenotype.

### Supplementary Figure 4

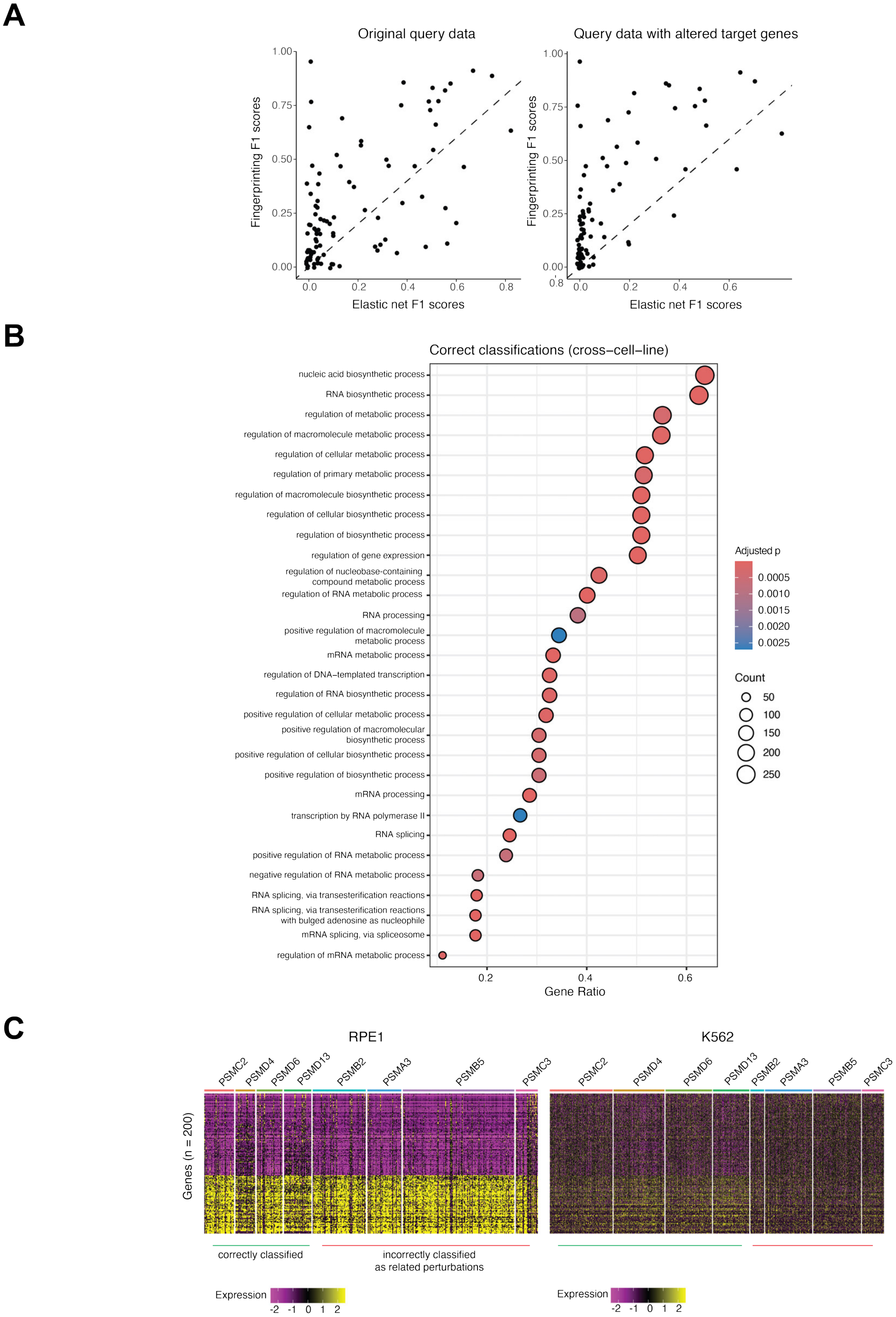

#### Supplementary Figure 4

(A) Scatterplots of F1 scores (as in Figure 4F) for each perturbation obtained under RNA fingerprinting versus elastic net, using the original query data (left) and with the target genes altered (right), to ensure that its reduction via CRISPRi does not singularly drive the match. F1 scores are shown for perturbations which were assigned at least once under both methods.
(B) Top gene ontology terms from enrichment analysis of the perturbations which were correctly classified across cell lines.
(C) Heatmap comparing gene expression for the same set of genes and perturbations in RPE1 cells (left) and K562 cells (right), with perturbations assigned correctly or incorrectly at the group-level indicated.

### Supplementary Figure 5

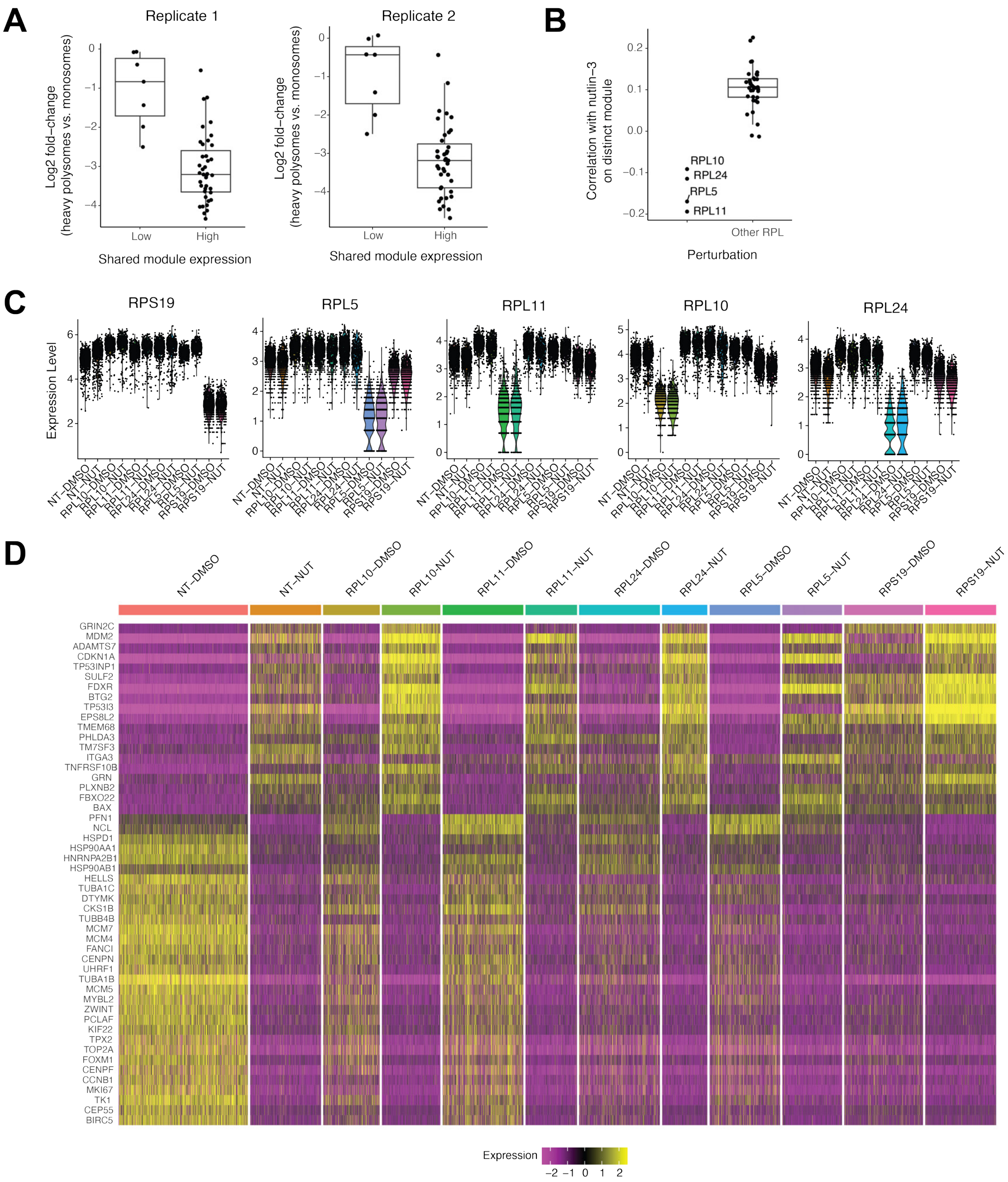

#### Supplementary Figure 5

(A) Log fold-changes of heavy polysomes to monosomes as reported by Nugent et al., 2024 for each RPL perturbation, split by their expression of our previously identified "shared RPL perturbation" module in the RPE1 data, hypothesized to correspond to the activation of ribosomal stress (Methods). Boxplots show data from two experimental replicates.
(B) Quantification of the gene expression patterns presented in Figure 5D-E. Boxplots show the median correlation (averaged across experimental replicates) between each RPL perturbation in the RPE1 data (as shown in Figure 5D) and the nutlin-3-treated cells in Figure 5E. Correlation is computed on genes comprising the distinct module in Figure 5D, for all RPL perturbations that successfully induced ribosomal stress.
(C) Validation of siRNA knockdown. Normalized expression of each targeted gene under each condition of the dsiRNA-mediated RPL depletion experiment.
(D) Heatmap of gene expression illustrating the expression of p53-responsive genes across cells from each of the selected RPL knockdowns, in both DMSO and nutlin-3a treated cells. Genes shown are the top differentially expressed genes between non-targeting cells treated with nutlin-3a and in DMSO.

### Supplementary Figure 6

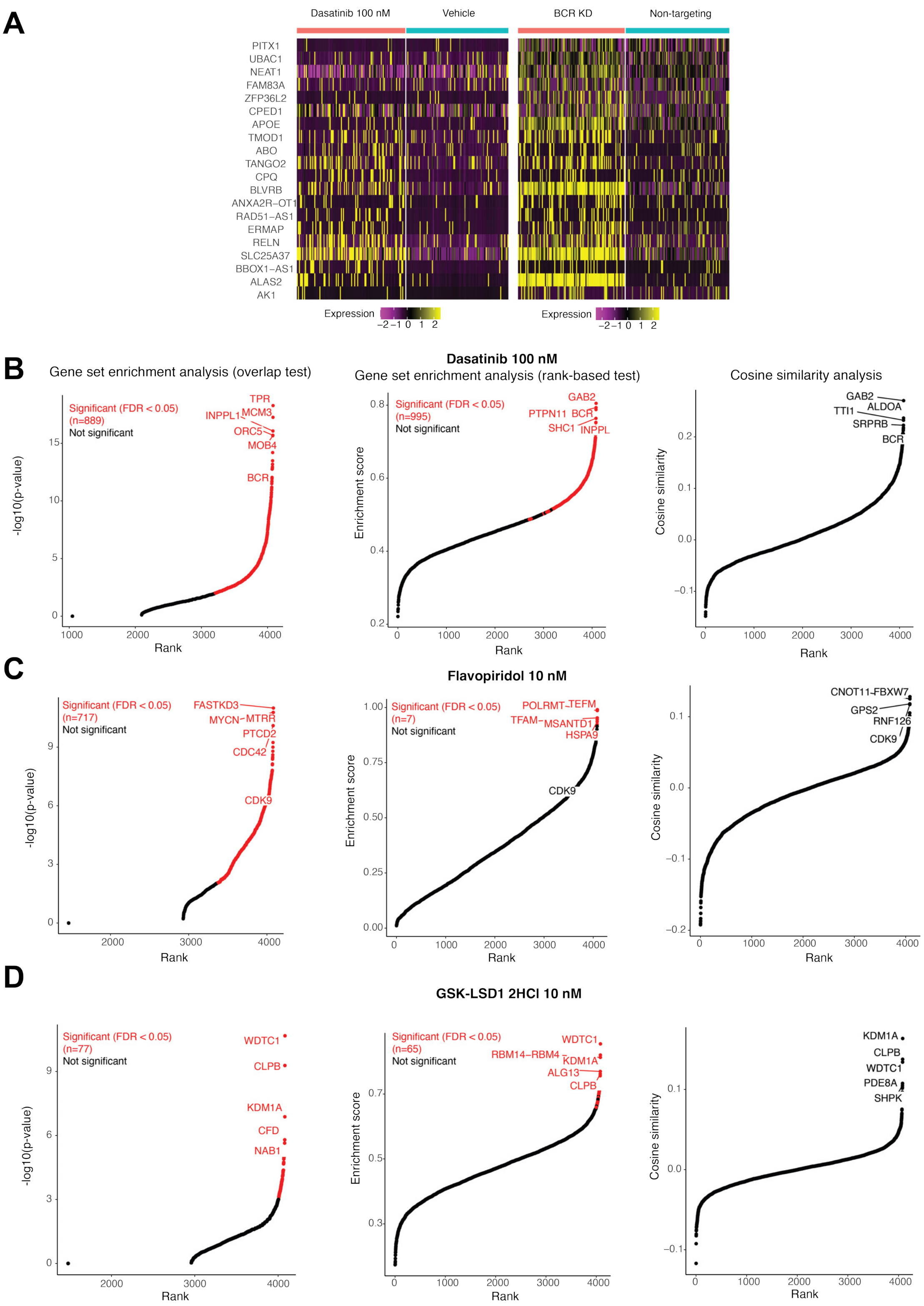

#### Supplementary Figure 6

(A) Heatmap showing the top genes driving the RNA fingerprinting match between Dasatinib 100 nM (sci-Plex) and BCR knockdown (GWPS). Genes included represent those that, when individually excluded from the model, resulted in the greatest reduction in Bayes factor (Methods). BCR knockdown cells are only shown if they were called perturbed by Mixscape analysis.
(B) Comparative approaches for assessing the target of Dasatinib 100 nM from GWPS (Methods). Left and middle plots show the overlap between differentially expressed genes computed based on the sci-Plex data (Dasatinib 100 nM vs. DMSO) and the GWPS data (each target gene vs. NT control). The true target (BCR) falls within a large list of significant putative matches, but is not the top hit. Right plot shows the cosine similarity between the scaled sci-Plex and GWPS expression profiles (Methods).
(C) As in (B), for Flavopiridol 10 nM.
(D) As in (B), for GSK-LSD1 2HCl 10 nM.

### Supplementary Figure 7

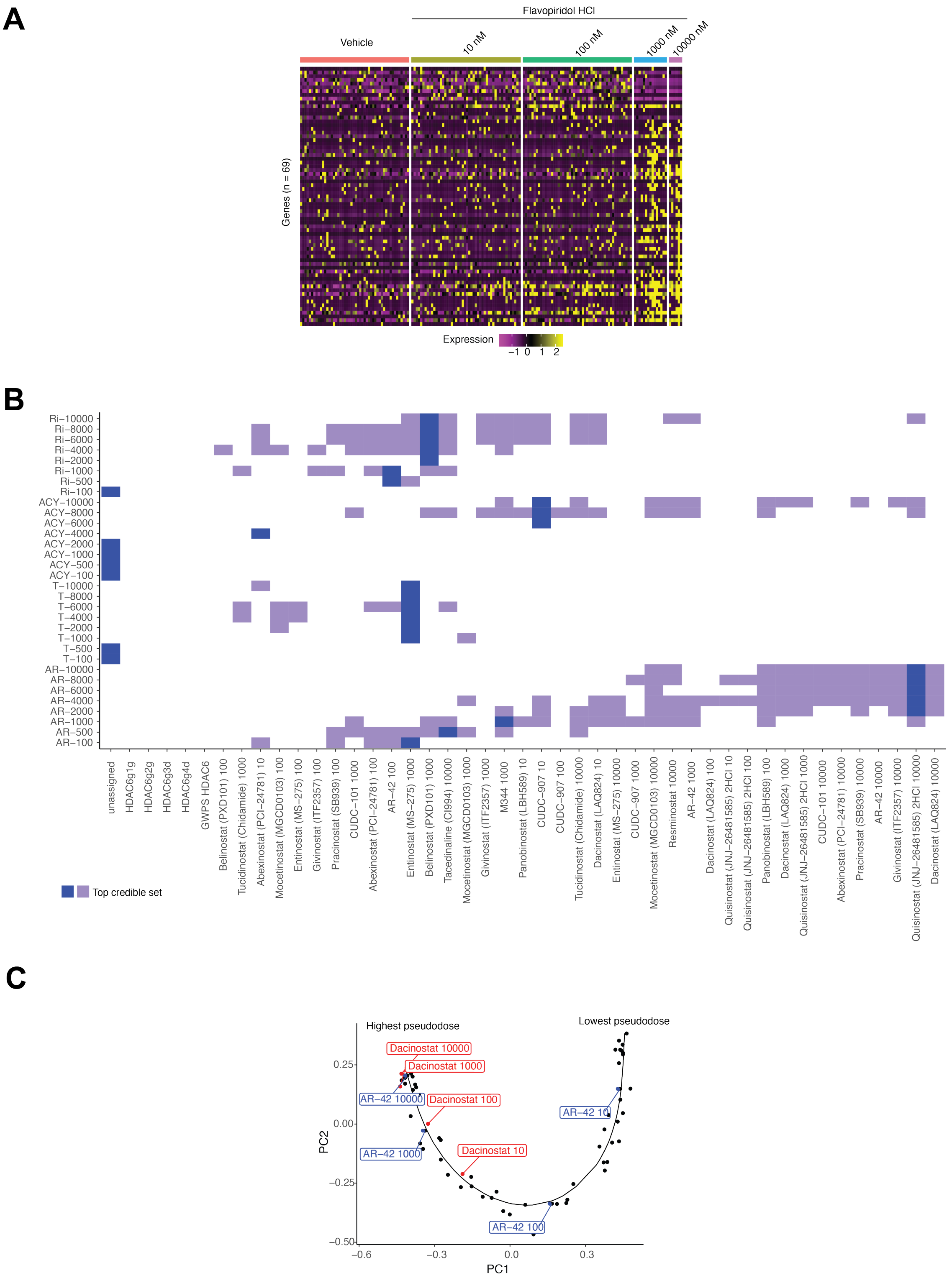

#### Supplementary Figure 7

(A) Heatmap showing gene expression profiles (Pearson residuals) of cells treated with Flavopiridol at increasing doses in the sci-Plex dataset.
(B) RNA Fingerprinting results for the FlexPlex experiment shown in Figure 6F. Shading indicates members of the top credible set for each drug-dose combination, darkest shade of blue indicates the top member. From this plot, we exclude two cases (ACY-738 500 nM and 2000 nM) where the coefficient of the top credible set was negative, as these were associated with low Bayes factors and were likely spurious matches.
(C) PCA plots of the fingerprints corresponding to HDAC inhibitors computed from the sci-Plex data. These fingerprints were overlaid with a fitted trajectory, corresponding to a "pseudodose" trajectory consistent with the sci-Plex manuscript. We used this trajectory to impute pseudodose levels for query drug profiles based on their RNA fingerprinting matches (Methods).

### Supplementary Figure 8

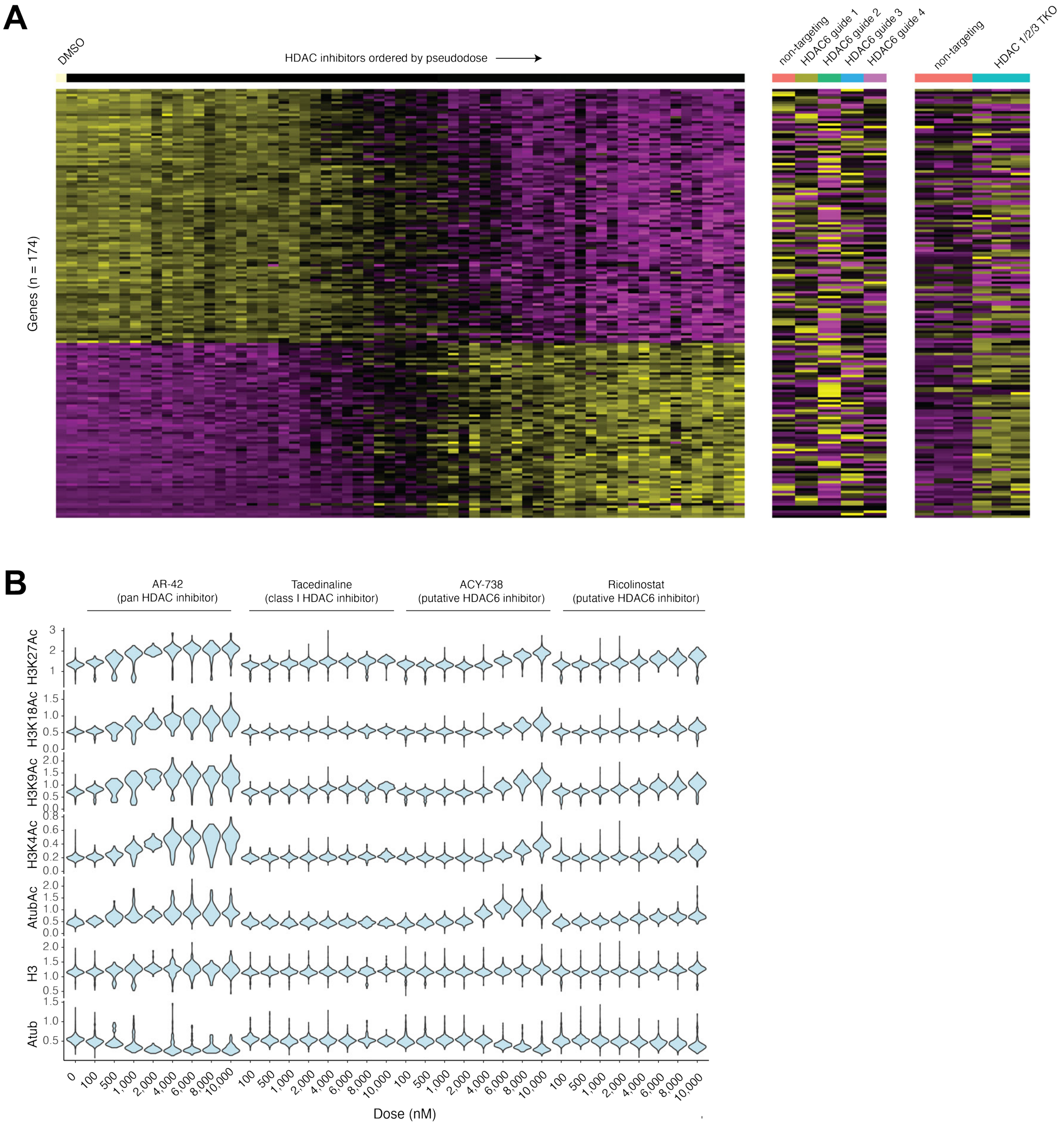

#### Supplementary Figure 8

(A) Gene expression heatmap for HDACi-dose combinations in the sci-Plex data. Each column represents a pseudobulked expression profile grouped by a drug-dose combination, and columns are ordered by pseudodose as computed in Supplementary Figure 7C. Trajectory-dependent genes are shown in the sci-Plex dataset (left), as well as pseudobulk profiles of HDAC6 CRISPRi data from this manuscript (middle) and HDAC 1/2/3 triple knock-out bulk RNA-seq data in HeLa cells from Li et al., 2023 (right).
(B) Normalized intracellular protein levels for each drug-dose combination as assessed in the FlexPlex experiment.

### Supplementary Figure 9

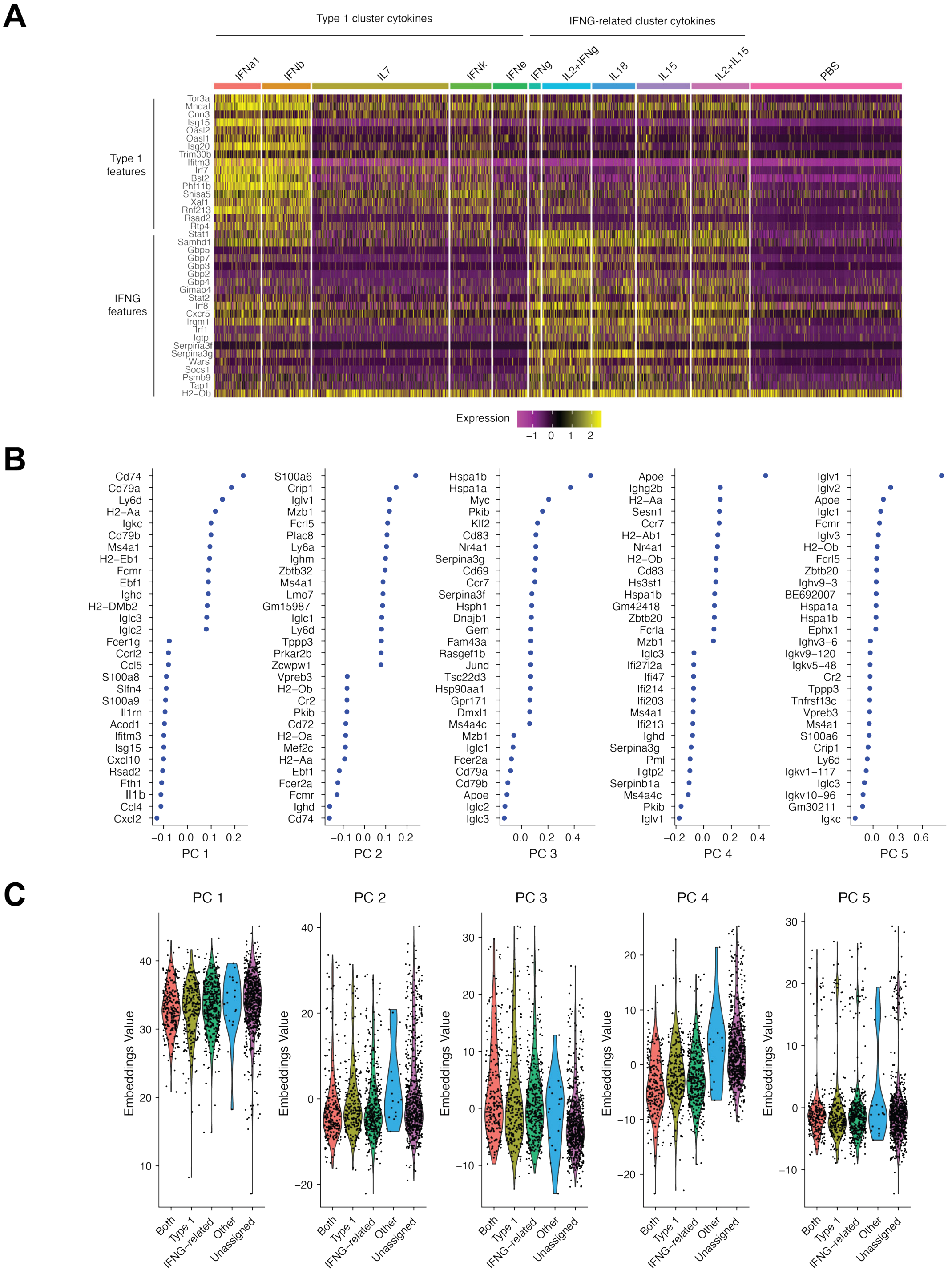

#### Supplementary Figure 9

(A) Gene expression programs associated with either Type 1-IFN or IFNG-related cytokine responses. Heatmap shows top genes associated with each cytokine cluster within B cells in the Immune Dictionary dataset. These genes are shown in Figure 7D to demonstrate heterogeneous activation within B cells responding to secondary flu infection.
(B) Features with top positive and negative loadings for the top five principal components from unsupervised analysis of the B cells shown in Figure 7E.
(C) Principal component scores for the top five principal components in (B). These unsupervised scores fail to separate the B cell subsets identified by RNA fingerprinting.
